# Regulatory-scale stripe analysis from single-cell Hi-C with scStripe

**DOI:** 10.64898/2026.09.16.751930

**Authors:** Ling Li, Yuchen Liu, Haishan Zhang, Wenjing Gao, Zeqin Lin, Dong Liu, Guangming Pan, Chi Yao, Dechao Tian

## Abstract

Chromatin stripes are a directional architectural feature of three-dimensional genome organization, but their analysis in single-cell Hi-C (scHi-C) at regulatory resolution is hindered by extreme sparsity. We developed scStripe, an imputation-free two-stage statistical framework that detects stripes from sparse cell-type–aggregated scHi-C maps and directly quantifies the resulting aggregate-defined stripes in raw individual-cell contact maps using per-cell stripe scores. Across three complementary benchmark settings, scStripe achieved the highest F1 scores and showed strong aggregate stripe enrichment, while the detected stripes recapitulated characteristic structural and regulatory features reported in bulk data. Per-cell stripe scores resolved recurrent configurations of the *EBF1*-anchored stripe, captured genome-wide variation associated with cell type, cell-cycle phase, and developmental stage, and identified positively associated gene–stripe pairs with pronounced cell-type specificity using matched scRNA-seq. Together, scStripe provides a practical, scalable framework for regulatory-scale stripe analysis across single cells without requiring single-cell enhancement or 3D reconstruction.

## Introduction

Chromatin stripes are directional, line-like submatrices of elevated interaction frequency in Hi-C maps that extend across contiguous genomic regions, indicative of one-sided loop extrusion [1, 2]. Stripes are associated with extended regulatory interactions that link promoters to distal cis-regulatory elements, including enhancers and silencers, at diverse genomic loci [2–7]. As a distinct feature of three-dimensional (3D) genome organization, stripes complement compartments, topologically associating domains (TADs), and chromatin loops [8–10]. Variation in stripe presence, strength, and directionality accompanies changes in chromatin folding and gene-regulatory landscapes across biological contexts including development, neurogenesis, and malignancy [3, 11–19]. Single-cell Hi-C (scHi-C), particularly when paired with matched transcriptomic measurements, provides an opportunity to examine stripe-associated variation across individual cells and relate this variation to gene expression in the same cells [20–28]. However, the extreme sparsity of scHi-C contact maps presents a major challenge for robust stripe detection and per-cell analysis at regulatory resolution [29].

Current stripe-calling approaches rely on high-resolution contact maps to resolve line-like interaction enrichment, yet raw scHi-C maps are extremely sparse at these resolutions, with a median missingness of approximately 99.94% within 2 Mb at 10 kb, complicating direct stripe detection in individual cells. Aggregating contacts from cells of the same type partially alleviates this sparsity and has enabled bulk stripe callers to be applied to cell-type–aggregated scHi-C maps [5, 21]. However, even after aggregation, these maps retain substantial sparsity, with approximately 31.41% of interactions missing within 2 Mb at 10 kb, and remain considerably sparser than the high-coverage bulk Hi-C maps for which existing stripe callers were developed [2, 30–34]. On these sparse aggregated maps, existing bulk stripe callers produce markedly different call sets and show method-dependent performance across complementary benchmarks and aggregate-enrichment measures (see our own analysis later). At the per-cell level, scHi-C enhancement methods improve the recovery of chromatin-contact patterns from sparse maps and provide a basis for downstream stripe analysis [35–38]. However, applying such methods genome-wide to every cell becomes increasingly memory-intensive at finer resolutions and larger cell numbers, with a recent benchmark reporting that several scHi-C workflows could not run at 200 kb for datasets of approximately 18,000 cells because of memory constraints, a resolution still substantially coarser than the 10 kb targeted here [38]. After aggregate-level stripe detection, prior studies have examined cell-to-cell variation by visualizing prominent stripes in selected cells [5] and characterizing multi-enhancer stripes at pre-selected marker-gene loci using scMicro-C coupled with high-resolution 3D reconstruction [21]. These studies provide valuable locus-specific insight, but extending such analyses genome-wide across large numbers of cells remains challenging, particularly when high-resolution 3D reconstruction is required. For example, Tensor-FLAMINGO requires more than 36 h to reconstruct 15 cells at 10 kb resolution [39]. Together, these advances and remaining limitations motivate a dedicated framework that couples robust stripe detection from sparse cell-type–aggregated scHi-C maps with direct, scalable quantification of aggregate-defined stripes across individual cells at regulatory resolution, without requiring single-cell enhancement or 3D reconstruction.

To address this need, we developed scStripe, an imputation-free two-stage framework for regulatory-scale stripe analysis of scHi-C data. In stage 1, scStripe detects stripes from sparse cell-type–aggregated scHi-C maps by combining random-matrix-theory–guided spectral selection and change-point detection with complementary statistical tests against matched backgrounds. In stage 2, scStripe quantifies the resulting aggregate-defined stripes directly in raw individual-cell contact maps using per-cell stripe scores that contrast pooled in-stripe contacts with matched flanking contacts. This ratio-based design enables direct per-cell quantification despite the extreme sparsity of raw scHi-C maps, without single-cell enhancement or 3D reconstruction. Across diverse datasets, scStripe detected stripes in nearly all evaluated cell types, achieved the highest F1 scores in three complementary benchmark settings, and showed strong aggregate stripe enrichment. The detected stripes also recapitulated characteristic structural and regulatory features of stripes reported in bulk data, including preferential localization in A compartments, frequent spanning of multiple TADs, and enrichment of CTCF and H3K4me3. At the per-cell level, scStripe resolved recurrent configurations of the *EBF1*-anchored stripe and captured multiple axes of biological variation through genome-wide per-cell stripe-score profiles. Integration with matched scRNA-seq further enabled genome-wide analysis of stripe–expression relationships in the same cells, identifying positively associated gene–stripe pairs with pronounced cell-type specificity. Together, scStripe provides a practical, scalable framework that connects aggregate-level stripe detection with direct per-cell quantification, enabling analysis of stripe variation and its relationship with matched gene expression across individual cells at regulatory resolution.

## Results

### Overview of scStripe

scStripe is an imputation-free two-stage framework for stripe analysis from sparse scHi-C contact maps at regulatory resolution, typically at 10 kb and at 5 kb for scMicro-C (Fig. 1). scStripe requires user-provided cell-type labels, which can be obtained from scHi-C-based clustering methods such as scHi-Cluster [35] or Fast-Higashi [40], or from external annotations such as matched single-cell RNA-seq profiles. In stage 1, contacts from cells of the same type are aggregated to generate normalized cell-type–aggregated scHi-C contact maps for stripe detection, whereas stage 2 quantifies the resulting aggregate-defined stripes directly in raw individual-cell contact maps without imputation (Fig. 1a). Aggregate-level detection is consistent with models in which stripes arise from recurrent directional interactions generated by one-sided loop extrusion or boundary stacking, such that aggregation can reinforce shared stripe-associated contact enrichment despite sparse observations in individual cells [2, 5]. For computational efficiency, chromosome-level aggregated maps are partitioned along the diagonal into overlapping 200×200-bin patches, following prior work [2, 31]. At 10 kb resolution, each patch spans 2 Mb along each genomic axis and encompasses 96.36% of the manually curated literature-reported stripes (Supplementary Table S1) and 99.41% of stripes called by Stripenn in bulk Hi-C maps, while reducing the matrix size processed relative to full-chromosome maps.

**Figure 1:**
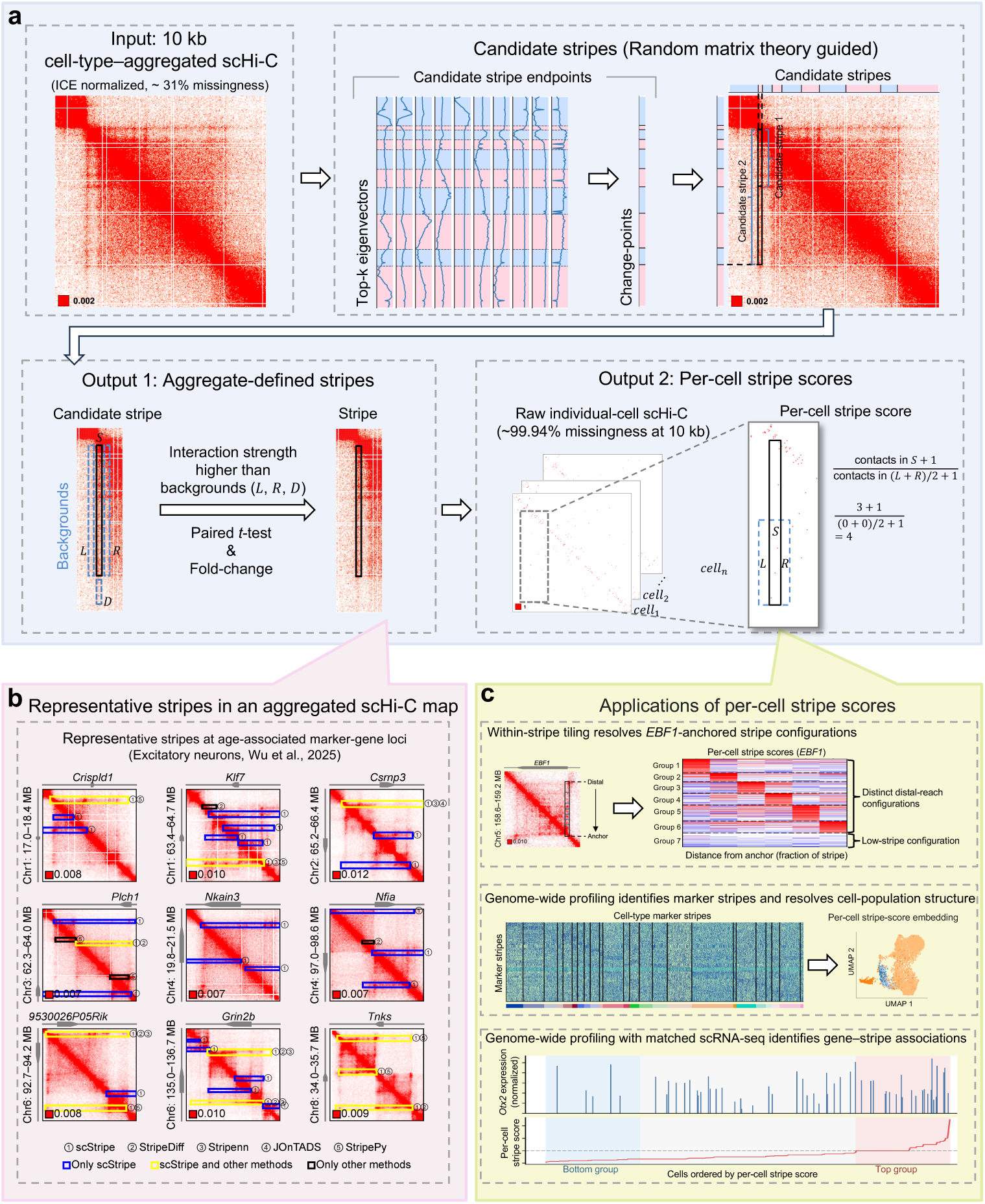
scStripe detects stripes from cell-type–aggregated scHi-C maps and quantifies them directly in individual cells. **a**, Overview of the two-stage scStripe framework. Cells of the same type are aggregated to generate normalized contact maps for stripe detection. Each contact-map patch is eigen-decomposed, and random-matrix-theory–guided spectral selection retains informative eigenvectors. Change-point detection identifies candidate endpoints. Endpoint triplets are assembled into directional stripe candidates, with two endpoints defining stripe width at the anchor and the third marking the distal end and thus stripe length. Candidates are evaluated using three one-sided tests and fold-change criteria. In stage 2, scStripe computes per-cell stripe scores for aggregate-defined stripes directly from raw individual-cell contact maps without imputation. The score contrasts pooled contacts in the stripe with those in matched flanks using the +1 pseudocount-stabilized ratio shown, enabling direct quantification despite the extreme sparsity of raw scHi-C maps. **b**, Representative stripes at nine age-associated marker-gene loci in the excitatory-neuron aggregated scHi-C map. For visualization, 3′-oriented stripes are shown in the upper triangle and 5′-oriented stripes in the lower triangle. Blue boxes denote stripes detected only by scStripe, yellow boxes those detected by scStripe and at least one alternative caller, and black boxes those detected only by alternative callers. **c**, Representative applications of per-cell stripe scores. Within-stripe tiling resolves recurrent configurations of the *EBF1*-anchored stripe. Genome-wide profiling identifies marker stripes and supports cell embedding from per-cell stripe scores. Matched scRNA-seq enables examination of stripe–expression relationships in the same cells, illustrated by an *Otx2* gene–stripe pair. Data shown in panels **b,c** are from the dscHi-C, scMicro-C and HiRES datasets [21, 22, 24], as indicated. Small boxes and numbers in the lower-left corners of contact maps indicate the upper limit of the color scale.

In stage 1, scStripe detects stripes from sparse, low-coverage aggregated scHi-C contact maps, which retain a median of 31.41% missingness within 2 Mb at 10 kb resolution. scStripe applies random-matrix-theory–guided spectral selection to retain informative eigenvectors, followed by change-point detection to identify candidate endpoints that are assembled into directional anchor-to-distal stripe candidates (Fig. 1a). Each candidate is evaluated using three complementary one-sided tests, comprising two tests of stripe enrichment relative to matched flanking backgrounds on either side and a distal-end test against a matched background beyond the detected endpoint, together with fold-change criteria for the flank comparisons. Candidates satisfying these criteria undergo de-duplication of redundant overlapping calls and constitute the final stripes called by scStripe. Representative stripes detected from the excitatory-neuron aggregated scHi-C map at age-associated marker-gene loci are shown in Fig. 1b. Component and parameter analyses further supported the stage-1 design. Increasing the change-point penalty reduced the number of change-points and detected stripes, while ASA score and stripiness increased gradually but only modestly, indicating a progressive trade-off between call-set size and stripe enrichment. A penalty of 0.1 was used as the default (Supplementary Fig. S1). The hypothesis-testing module retained stripes with consistent ASA enrichment that would be removed by stripiness-based filtering (Supplementary Fig. S2), while stepwise analysis showed that the three tests progressively refined the candidate set (Supplementary Fig. S3). Across alternative fold-change and significance thresholds, stripe calls remained broadly consistent and maintained concordance with lineage-related bulk Hi-C (Supplementary Fig. S4), while ASA scores and CTCF and H3K4me3 enrichment patterns remained similar across parameter settings (Supplementary Fig. S5).

In stage 2, scStripe quantifies each aggregate-defined stripe directly in raw individual-cell contact maps without imputation or 3D reconstruction (Fig. 1a). These raw maps are extremely sparse, with approximately 99.94% missingness within 2 Mb at 10 kb resolution. For each cell, the per-cell stripe score pools contacts across the stripe and its matched flanks and quantifies stripe enrichment relative to the combined flanking contacts using a +1 pseudocount. Because the score is defined as a ratio between pooled contacts in the stripe and matched flanks, it is expected to provide a stable measure of relative stripe enrichment despite the extreme sparsity of raw scHi-C contact maps. This expected robustness is evaluated empirically by contact-downsampling analyses described below. The score operates in two complementary settings. Genome-wide profiling assigns one score to each stripe in every cell, using the distal 50% of the stripe body by default to reduce the influence of near-diagonal enrichment, whereas within-stripe tiling computes multiple segment-level scores across a selected stripe to resolve internal heterogeneity. The resulting stripe-by-cell and tiled subregion-by-cell matrices provide quantitative representations of stripe variation across individual cells and support downstream analyses including marker-stripe identification, cell embedding, and integration with co-assayed scRNA-seq profiles (Fig. 1c).

To evaluate scStripe across controlled levels of scHi-C sparsity, we independently downsampled contacts from excitatory neurons to 10–75% of the full sequencing depth and assessed both stages of the framework, as detailed in Supplementary Methods. For stage 1, stripe-calling concordance between two non-overlapping cell groups remained stable at sampling depths of 25% and above, while the resulting stripes retained similar CTCF and H3K4me3 enrichment patterns across depths (Supplementary Fig. S6). For stage 2, per-cell stripe-score profiles showed increasing agreement with the full-depth data as sequencing depth increased, while cell–cell distance structure and stripe-score variability showed similar depth-dependent agreement (Supplementary Fig. S7). To assess potential self-inclusion effects, cells were divided into two complementary, non-overlapping groups, and stripes identified from each group were scored either in the same group used for stripe detection (within-group) or in the complementary group, whose cells had not contributed to stripe detection (held-out). Similar results under within-group and held-out scoring suggested limited sensitivity of per-cell stripe-score estimation to self-inclusion (Supplementary Fig. S7). Together, these analyses support the robustness of both aggregate-level stripe detection and direct per-cell stripe quantification across varying levels of scHi-C sparsity and sequencing depth.

### Systematic benchmarking of stripe detection in aggregated scHi-C contact maps

To systematically evaluate stripe detection from sparse aggregated scHi-C contact maps, we benchmarked scStripe against six alternative stripe callers, including Stripenn [31], StripeDiff [32], JOnTADS [33], StripePy [34], StripeCaller and Zebra [2, 31]. The benchmark comprised nine single-cell chromatin conformation datasets from seven published studies, collectively covering 94 annotated cell types across multiple species and sequencing depths [21–27]. These datasets were generated using diverse single-cell chromatin conformation assays, including Dip-C, HiRES, GAGE-seq, LiMCA, dscHi-C, ChAIR and scMicro-C. For the large-scale benchmark, scStripe, Stripenn, StripeDiff, JOnTADS, StripePy and StripeCaller were applied at 10 kb resolution to the same ICE-normalized aggregated scHi-C contact maps. Because Zebra requires a distinct preprocessing workflow, it was included in focused comparisons rather than the full cross-cell-type benchmark.

The breadth of stripe detection across cell types was first assessed. Among the 94 cell types, scStripe detected at least one stripe in 92 cell types, with StripeDiff detecting stripes in a similar number of cell types (91), whereas the remaining callers detected stripes in substantially fewer cell types (54–72; Supplementary Fig. S8a). For quantitative comparability, subsequent cross-cell-type comparisons were restricted to the 42 cell types across eight datasets in which all six methods included in the large-scale benchmark detected at least one stripe.

The number and overlap of detected stripes were next compared. At nine representative age-associated genomic loci in excitatory neurons, scStripe identified 33 stripes, including 22 detected uniquely by scStripe and 11 also detected by at least one alternative caller, whereas 4 stripes were detected only by other callers (Fig. 1b). Genome-wide, scStripe detected 5,697 stripes in excitatory neurons (Fig. 2a). This call-set size is of the same order of magnitude as the approximately 4,000 stripes reported in high-coverage bulk Hi-C maps of activated B cells [2]. StripeCaller and Zebra also produced large call sets, with StripeCaller detecting more stripes than scStripe and Zebra detecting a similar number. In contrast, StripeDiff, Stripenn, JOnTADS and StripePy each detected no more than 982 stripes, such that scStripe detected at least 5.8 times as many stripes as each of these methods (Fig. 2a). Of the merged stripes detected by the six alternative callers, 59.8% overlapped scStripe calls (Fig. 2a).

**Figure 2:**
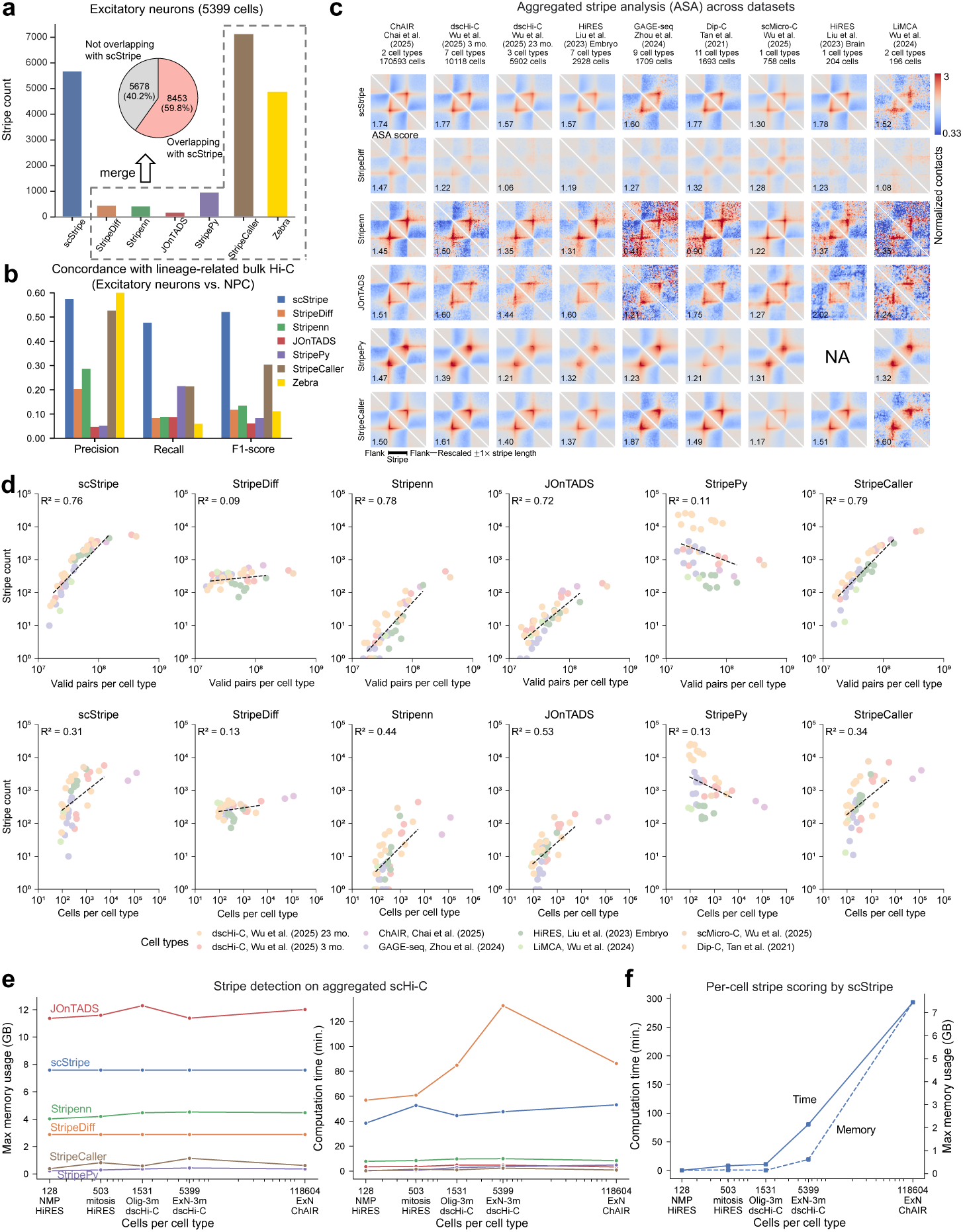
scStripe detects stripes broadly from aggregated scHi-C maps while retaining strong support across complementary benchmarks and aggregate stripe enrichment. **a**, Stripe counts for scStripe and six alternative callers in the aggregated scHi-C map of excitatory neurons (5,399 cells). The inset pie chart shows the fraction of merged stripes detected by the six alternative callers that overlap scStripe calls. **b**, Concordance between stripes detected from the excitatory-neuron aggregated scHi-C map and method-specific stripe sets called by the same method from lineage-related NPC bulk Hi-C. **c**, Aggregated stripe analysis (ASA) of stripes detected by scStripe and alternative callers across nine datasets. For each dataset, stripe-centered windows were rescaled to *±*1*×* stripe length and averaged across cell types in which the corresponding method detected at least one stripe. The ASA score, shown in the lower-left corner of each heatmap, is defined as the mean normalized contact intensity within the central stripe submatrix divided by that in the upper-right flanking submatrix. NA indicates dataset–method combinations for which no cell type met the requirement of at least one detected stripe. **d**, Relationship between stripe count and sequencing depth (*top*) or number of aggregated cells (*bottom*) across cell types. Dashed lines show linear regression fits. **e**, Peak memory usage (*left*) and runtime (*right*) for stripe detection on chromosome 1 across cell types containing 128–118,604 cells. **f**, Runtime and peak memory usage for scStripe per-cell stripe scoring across all chromosomes for cell types containing 128–118,604 cells. All analyses were performed at 10 kb resolution. Panels **a** and **b** use excitatory-neuron aggregated scHi-C and lineage-related NPC bulk Hi-C data from [22, 41], whereas panels **c–f** use datasets from [21–27]. Zebra was included only in the focused comparisons in **a** and **b**. For panels **d–f**, axes are presented on a log10 scale (d: both axes; e,f: x-axis only).

Extending the comparison across the 42 cell types showed substantial method-dependent variation in stripe counts (Supplementary Fig. S8b–d). scStripe detected more stripes than Stripenn, StripeDiff and JOnTADS, with median fold increases of 50.4×, 2.89× and 41.17×, respectively. In contrast, StripePy generally detected more stripes than scStripe, with a median scStripe-to-StripePy ratio of 0.22× (Supplementary Fig. S8b,c). scStripe and StripeCaller produced broadly comparable numbers of stripes across cell types (Supplementary Fig. S8b,c). Across the 42 cell types, a median of 41.6% of Stripenn stripes overlapped scStripe calls (Supplementary Fig. S8d).

Stripe detection was next evaluated using three complementary benchmark settings. First, concordance with lineage-related bulk Hi-C was assessed for stripes detected in the excitatory-neuron aggregated scHi-C map. Each method was compared with a method-specific reference set defined by stripes called by the same method in NPC bulk Hi-C [41]. scStripe achieved the highest F1 score of 0.52, compared with 0.30 for StripeCaller, the second-best method (Fig. 2b). Second, stripe detection accuracy was evaluated under controlled simulation using SimuStripe, in which the embedded stripes are predefined [32]. This benchmark was limited to scStripe, StripeDiff and JOnTADS, which can operate directly on the simulated contact matrices. scStripe again achieved the highest F1 score of 0.65, compared with 0.54 for StripeDiff, the second-best method (Supplementary Fig. S9a). Third, performance was evaluated against a manually curated set of 55 literature-reported stripes from seven bulk Hi-C studies (Methods; Supplementary Table S1). On the subset of 51 stripes for which all seven callers could be evaluated on the corresponding high-coverage bulk Hi-C maps, scStripe achieved the highest F1 score of 0.59, narrowly exceeding Stripenn at 0.57 (Supplementary Fig. S9b,c). Stripenn achieved higher precision than scStripe (0.76 versus 0.68), whereas scStripe achieved higher recall (0.53 versus 0.46). Across the three complementary benchmark settings, scStripe achieved the highest F1 score within each setting, whereas the second-best method differed among settings.

Next, aggregate stripe signal was assessed using aggregated stripe analysis (ASA), which quantifies contact enrichment within stripe submatrices relative to matched flanking regions. scStripe produced consistently continuous anchor-to-distal enrichment patterns in ASA heatmaps across datasets (Fig. 2c). Across the 42 cell types, scStripe achieved the highest median ASA score of 1.67, compared with 1.61 for JOnTADS, the second-highest method (Supplementary Fig. S8e). These results show that scStripe-detected stripes retain strong aggregate anchor-to-distal interaction enrichment across diverse cell types and datasets.

Finally, the relationship between data size, stripe detection and computational requirements was evaluated. Stripe counts from scStripe and several alternative callers increased with sequencing depth and, to a lesser extent, with the number of aggregated cells, whereas these associations varied substantially among methods (Fig. 2d). For stripe detection on chromosome 1, scStripe showed runtime and peak memory requirements comparable to those of alternative stripe callers under single-threaded execution, requiring approximately 1 h and 8 GB of peak memory for the largest evaluated cell type containing 118,604 cells (Fig. 2e). For scStripe per-cell stripe scoring across all chromosomes, runtime increased with cell number while peak memory usage remained stable. For the largest evaluated cell type, per-cell stripe scoring required approximately 5 h and 8 GB of peak memory, remaining computationally feasible at the largest evaluated data size (Fig. 2f).

Together, these systematic benchmarks across diverse scHi-C datasets showed that scStripe detects stripes across nearly all evaluated cell types from sparse aggregated contact maps, achieves the highest F1 scores across complementary benchmark settings, retains strong aggregate stripe enrichment, and remains computationally feasible for large cell populations.

### Stripes detected from aggregated scHi-C maps show characteristic structural and regulatory chromatin features

Bulk Hi-C studies have linked chromatin stripes to A/B compartmentalization and TAD organization [2, 31]. To determine whether these structural relationships are retained in stripes detected from sparse aggregated scHi-C maps, we examined their associations with A/B compartments and TADs called from the same aggregated contact maps used for stripe detection (Methods). To use a common set of cell types across both analyses, the 11 Dip-C-derived cell types [27] were excluded because TADs could not be called using the standard cooltools workflow [42], leaving 31 cell types across seven datasets for analysis.

To examine stripe–compartment relationships, stripes were classified as residing entirely within A compartments (only A), entirely within B compartments (only B), or spanning both compartments (mixed; Fig. 3a). For this analysis, up to 200 stripes per method and cell type were selected to reduce differences in call-set size across methods. Stripes were ranked using Stripenn-derived *P* values, with all stripes retained when fewer than 200 were detected. In excitatory neurons, 48% of scStripe-detected stripes were located only in A compartments, compared with 15% located only in B compartments (Fig. 3b). A similar A-compartment preference was observed for most alternative callers, including Zebra, in this focused comparison. Across the 31 cell types, scStripe-detected stripes remained preferentially located in A compartments, with median proportions of 56.0% for only A stripes and 13.5% for only B stripes (Fig. 3d). Stripenn, JOnTADS, StripePy and StripeCaller showed similar A-compartment enrichment (median only-A proportions, 51.5–71.76%), whereas StripeDiff showed no comparable preference (27.0% only A versus 30.0% only B). By contrast, TADs were distributed more evenly between A and B compartments, with median proportions of 45.3% and 44.4%, respectively.

**Figure 3:**
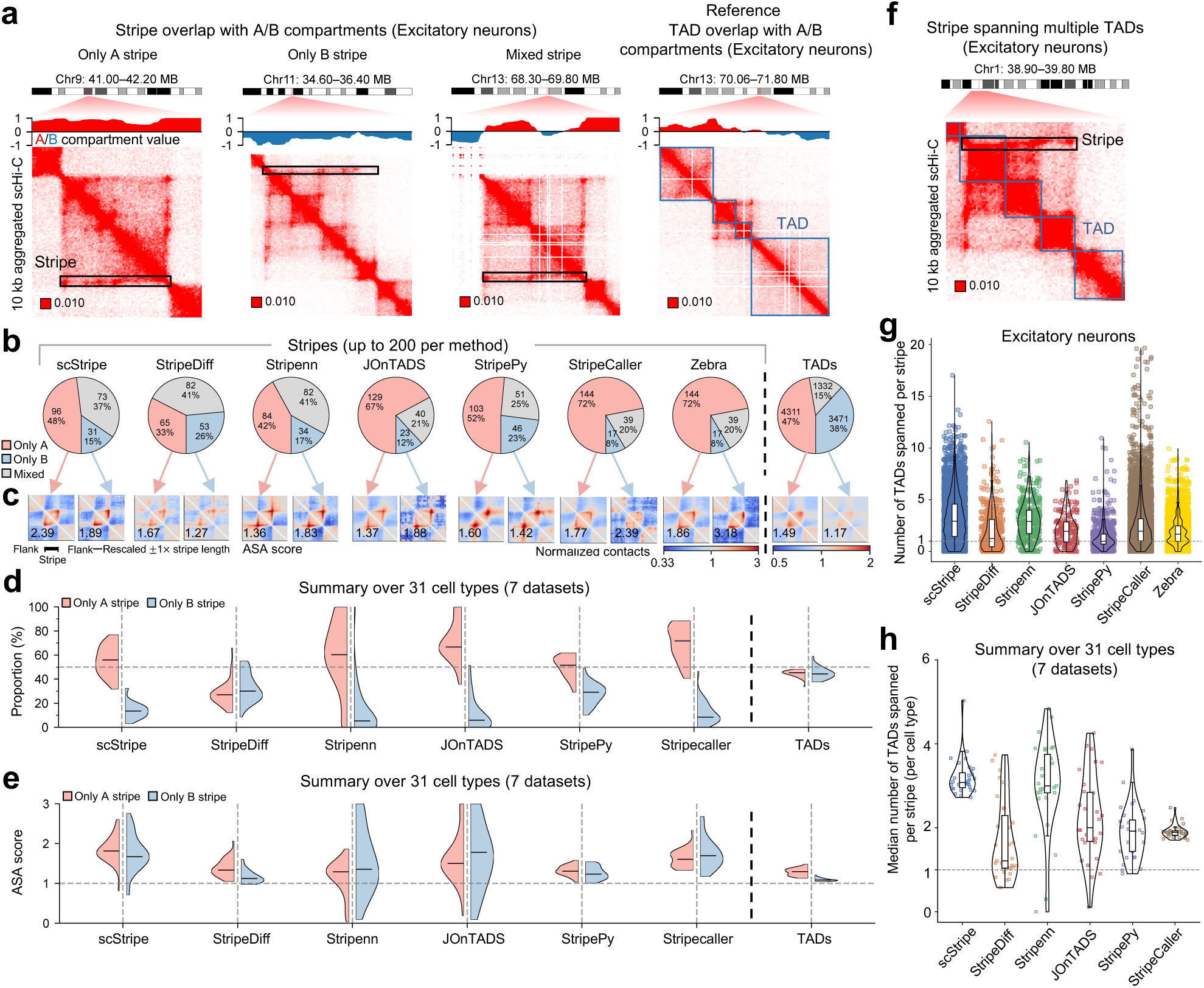
scStripe-detected stripes are preferentially located in A compartments and frequently span multiple TADs. **a**, Representative excitatory-neuron aggregated scHi-C contact maps showing scStripe-detected stripes located entirely within A compartments (only A), entirely within B compartments (only B), or spanning both compartments (mixed). Representative TADs located entirely within A or B compartments are shown as structural references. **b**, Proportions of only A, only B and mixed stripes across stripe callers in excitatory neurons, with TADs shown as a structural reference. **c**, ASA heatmaps for only A and only B stripes in excitatory neurons, with TADs included as a structural reference. ASA scores are shown in the lower-left corner of each heatmap. **d**, Distributions of the proportions of only A and only B stripes across 31 cell types from seven datasets, with TADs shown as a structural reference. The horizontal dashed line marks 50%. **e**, Compartment-stratified ASA scores across the same 31 cell types. The horizontal dashed line marks the ASA baseline of 1. Only compartment-specific stripe or TAD sets containing at least one feature were included in ASA. **f**, Representative scStripe-detected stripe spanning three consecutive TADs in excitatory neurons. **g**, Distributions of the number of TADs spanned per stripe for all stripes detected by each method in excitatory neurons. **h**, Distributions of the cell-type-level median number of TADs spanned per stripe across 31 cell types from seven datasets. The horizontal dashed line marks one TAD. For the compartment analyses in **b**–**e**, up to 200 stripes per method and cell type were selected by ranking calls using Stripenn-derived *P* values, with all detected stripes retained when fewer than 200 were available. Panels **f**–**h** used all stripes detected by each method. Stripes and TADs were analyzed at 10 kb resolution and A/B compartments at 50 kb resolution, all derived from the same aggregated scHi-C maps. Zebra was evaluated only in the focused excitatory-neuron comparisons in **b**, **c** and **g**.

To determine whether stripes in both compartment classes retained stripe-associated interaction enrichment, ASA was calculated separately for only A and only B stripes, with TADs included as a structural reference. In excitatory neurons, scStripe-detected stripes showed ASA scores of 2.39 in A compartments and 1.89 in B compartments, both exceeding the baseline of 1 and the corresponding TAD scores of 1.49 and 1.17 (Fig. 3c). Elevated ASA was also observed for both compartment classes across alternative callers. Across the 31 cell types, scStripe showed median ASA scores of 1.80 for only A stripes and 1.74 for only B stripes, whereas TAD ASA scores were distributed closer to 1 (Fig. 3e). Thus, although scStripe-detected stripes were preferentially located in A compartments, those residing in B compartments also retained clear aggregate stripe enrichment.

We next characterized the relationship between stripes and TADs by quantifying the number of TADs spanned by each stripe, using all stripes identified by each method. Consistent with observations from bulk Hi-C [2], scStripe-detected stripes frequently spanned multiple TADs, as illustrated by a representative stripe spanning three consecutive TADs in excitatory neurons (Fig. 3f). In excitatory neurons, scStripe-detected stripes spanned a median of approximately 3 TADs per stripe (Fig. 3g). Across the 31 cell types, the median number of TADs spanned per stripe was approximately 3 for both scStripe and Stripenn, compared with 1–2 for StripeDiff, JOnTADS, StripePy and StripeCaller (Fig. 3h).

To further characterize the regulatory chromatin context of the detected stripes, ChIP-seq data for the architectural protein CTCF and the active promoter-associated histone mark H3K4me3 from lineage-related NPCs were examined in relation to stripes detected from the excitatory-neuron aggregated scHi-C map. TADs were classified as stripy when they overlapped at least one detected stripe and as non-stripy otherwise. For scStripe and several alternative callers, the boundaries of stripy TADs showed higher CTCF and H3K4me3 enrichment than the corresponding boundaries of non-stripy TADs (Supplementary Fig. S10a–c), consistent with bulk-data observations linking stripy TADs to architectural-protein occupancy and active chromatin [31]. When ChIP-seq signals were aligned to stripe coordinates, both CTCF and H3K4me3 showed similar enrichment patterns along stripe coordinates, with enrichment within stripe regions and increased signals at stripe anchors and toward distal ends for scStripe and several alternative callers (Supplementary Fig. S10d,e). These patterns are consistent with bulk-data observations of architectural-protein enrichment at stripe anchors and active chromatin enrichment within stripe-associated regions [31].

Together, these analyses show that stripes detected from aggregated scHi-C maps exhibit characteristic structural and regulatory chromatin features reported in bulk data, including preferential localization in A compartments, frequent spanning of multiple TADs and enrichment of CTCF and H3K4me3. These results support the use of aggregate-defined stripes for subsequent per-cell quantification.

### Per-cell stripe scores resolve recurrent configurations of the *EBF1*-anchored stripe

A recent scMicro-C study identified an approximately 500 kb promoter-anchored stripe at the *EBF1* locus and revealed heterogeneous promoter–enhancer interactions across individual cells using reconstructed single-cell 3D genome structures [21]. To assess whether scStripe-derived per-cell stripe scores capture this heterogeneity directly from single-cell contact maps, the same scMicro-C dataset was reanalyzed. scStripe reproduced the reported *EBF1* stripe in a 5 kb aggregated scMicro-C contact map generated from 758 GM12878 cells (Fig. 4a). Because a single score for the entire stripe would summarize overall stripe strength but not variation in distal reach, the stripe body was partitioned along the distal-to-anchor axis into six enhancer-guided tiled subregions. A separate per-cell stripe score was calculated for each subregion, yielding six tiled stripe scores that together represent the local stripe profile of each cell.

**Figure 4:**
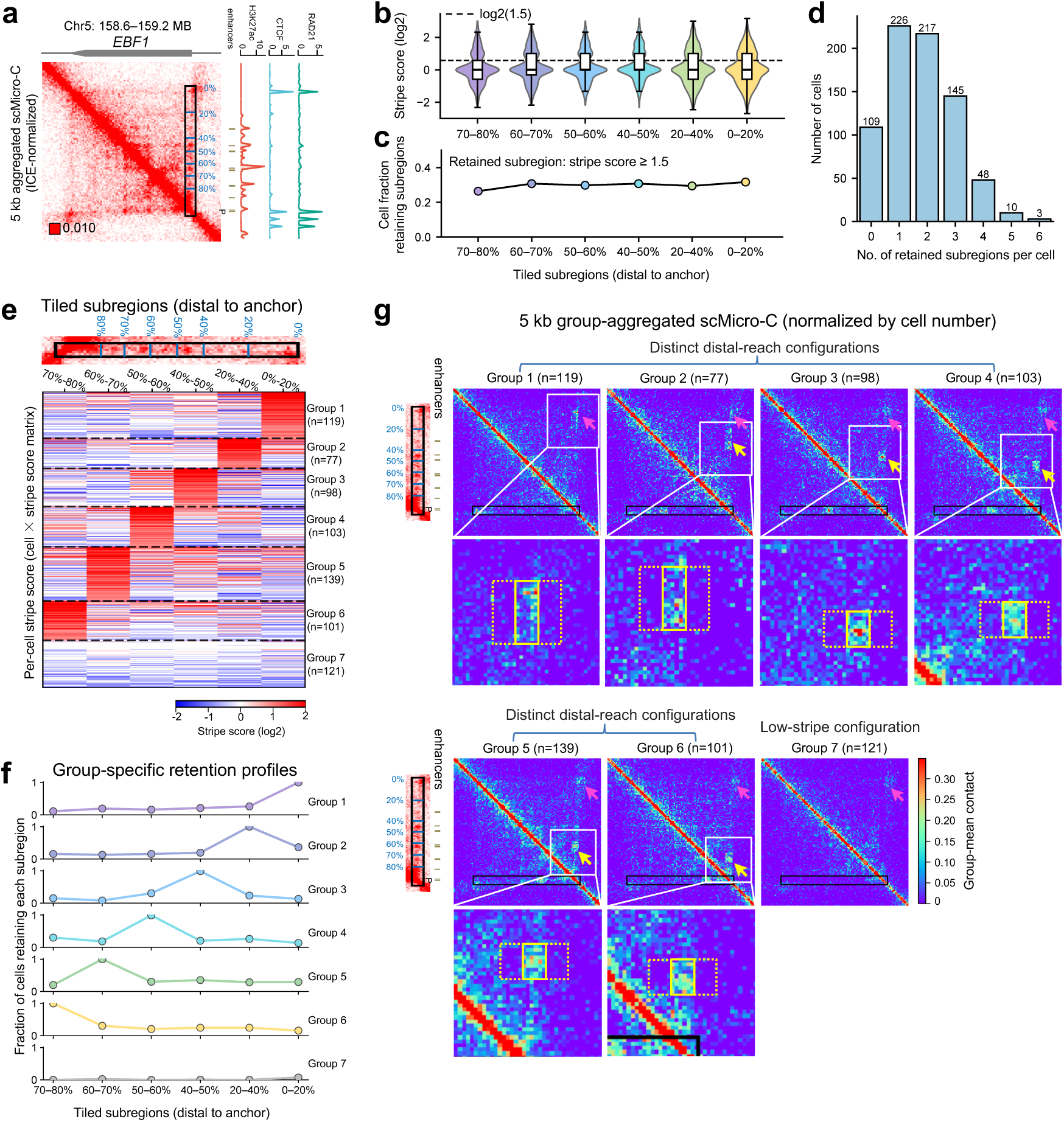
Per-cell stripe scores resolve recurrent configurations of the *EBF1*-anchored stripe. **a**, scStripe-detected promoter-anchored stripe spanning approximately 500 kb at the *EBF1* locus in a 5 kb aggregated scMicro-C contact map generated from 758 GM12878 cells. The stripe extends across seven ABC-predicted enhancers (E1–E7) and is shown together with CTCF, RAD21 and H3K27ac ChIP-seq tracks. This stripe was previously reported in the scMicro-C study [21]. **b**, Violin plots of log_2_ per-cell stripe scores across six enhancer-guided tiled subregions ordered from distal to anchor. The dashed line marks log_2_(1.5). **c**, Fraction of cells retaining each tiled subregion, with retention defined as stripe score *≥* 1.5. **d**, Distribution of the number of retained subregions per cell. **e**, Heatmap of the six tiled per-cell stripe scores after hierarchical clustering, resolving seven groups with distinct score patterns across the tiled subregions. **f**, Group-specific fractions of cells retaining each tiled subregion at a stripe-score threshold of 1.5. **g**, Group-aggregated 5 kb contact maps normalized by cell number, showing distinct distal-reach configurations across Groups 1–6 and a low-stripe configuration in Group 7. Zoomed regions highlight group-specific stripe patterns. Yellow arrows indicate selected enhancer-associated contacts, solid yellow boxes mark stripe subregions, dotted yellow boxes mark flanking regions, black boxes indicate the stripe detected in the aggregated map of all 758 cells and pink arrows indicate CTCF-associated loops. All analyses were performed at 5 kb resolution. CTCF, RAD21 and H3K27ac tracks were obtained from 4DN; ABC-predicted enhancer annotations and scMicro-C data are from Wu et al. [21].

The six tiled stripe scores varied substantially among the 758 GM12878 cells, revealing pronounced cell-to-cell heterogeneity across the *EBF1*-anchored stripe (Fig. 4b–d; Supplementary Fig. S11a). Using a stripe-score threshold of 1.5 to define retention of a tiled subregion, individual subregions were retained in only 26.39–31.66% of cells (Fig. 4c). Most cells retained only one or two subregions, with 29.82% retaining one and 28.63% retaining two, whereas only 8.05% retained four or more subregions (Fig. 4d). These distributions were qualitatively stable when the retention threshold was varied from 1.1 to 2.0 (Supplementary Fig. S11b,c). Thus, although the aggregated contact map shows a prominent *EBF1*-anchored stripe extending across multiple enhancer-containing subregions, the corresponding per-cell stripe profiles are highly heterogeneous, consistent with the variable distal enhancer engagement reported in the scMicro-C study [21].

Cells were next grouped according to their six tiled stripe scores to determine whether the observed heterogeneity could be organized into recurrent stripe configurations. Hierarchical clustering resolved seven groups containing 77–139 cells each (Fig. 4e). Groups 1–6 showed distinct high-score patterns across the tiled subregions, whereas Group 7 showed uniformly low stripe scores across all six subregions (mean ≤ 0.89; Fig. 4e,f; Supplementary Fig. S11e). The group-specific retention profiles remained qualitatively stable when the retention threshold was varied from 1.1 to 2.0 (Supplementary

Fig. S11d). The same organization was evident in group-aggregated 5 kb contact maps, which showed distinct promoter-anchored stripe patterns across Groups 1–6 and no comparable stripe pattern in Group 7 (Fig. 4g). These concordant score and contact-map patterns support six recurrent promoter-anchored stripe configurations with distinct dominant distal subregions, consistent with the heterogeneous EBF1 promoter–enhancer interactions reported in the scMicro-C study [21]. By contrast, Group 7 showed uniformly low stripe scores and no comparable stripe in the group-aggregated contact map, representing an additional low-stripe configuration not described in that study.

Together, these analyses show that scStripe resolves heterogeneous *EBF1*-anchored stripe configurations directly from raw single-cell contact maps, without the single-cell 3D genome reconstruction used in the scMicro-C study [21]. More broadly, per-cell stripe scores provide a quantitative representation of stripe variation for direct comparisons across cells and downstream associations with matched scRNA-seq profiles.

### Genome-wide per-cell stripe scores capture multiple axes of biological variation

We next asked whether per-cell stripe scores capture biological variation across cells beyond individual loci. Unlike the within-stripe tiling mode used for the *EBF1*-anchored stripe, genome-wide profiling assigns one score to each detected stripe in every cell, using the distal 50% of the stripe body relative to matched flanking regions. In the co-assayed HiRES dataset [24], the resulting stripe-by-cell matrix contained cell-type-specific marker stripes (Fig. 5a). An embedding based solely on genome-wide per-cell stripe scores separated the four major cell populations and also reflected variation associated with cell-cycle phase and, within embryonic lineages, developmental stage (Fig. 5b), consistent with patterns reported using scHiCluster-enhanced scHi-C in the original HiRES study [24]. These embedding patterns were qualitatively stable across alternative analysis settings (Supplementary Fig. S12). Similar cell-type-specific marker stripes and separation of annotated cell types were observed in an independent dscHi-C dataset [22] (Supplementary Fig. S13).

**Figure 5:**
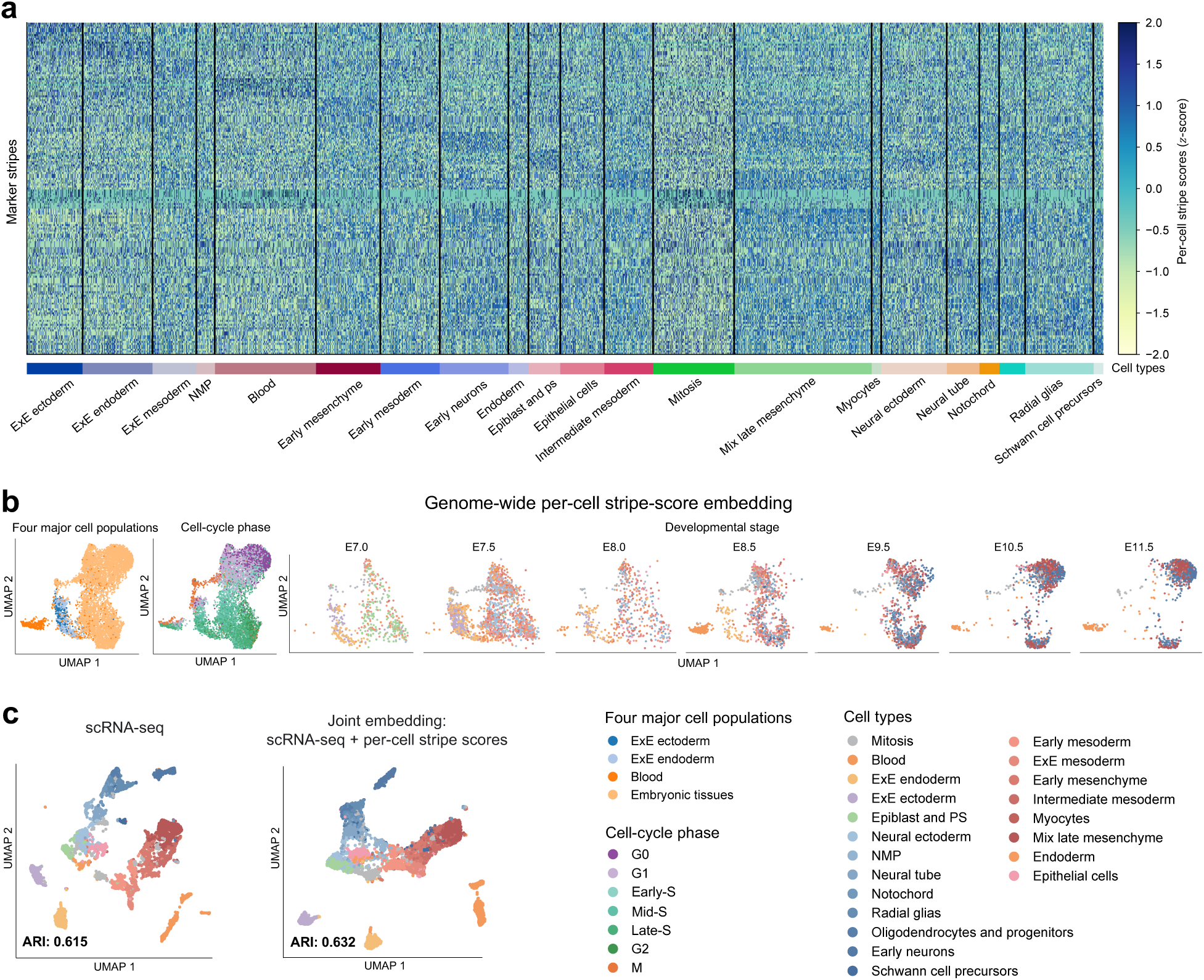
Genome-wide per-cell stripe scores capture multiple axes of biological variation. **a**, Heatmap of *z*-scored per-cell stripe scores for cell-type marker stripes, with cells grouped by annotated cell type. **b**, UMAP embedding based solely on genome-wide per-cell stripe scores. The same embedding is shown by four major cell populations and cell-cycle phase, followed by stage-specific views spanning E7.0–E11.5, with cells colored by annotated cell type. **c**, UMAP embeddings based on scRNA-seq alone (*left*) and joint embedding of scRNA-seq and genome-wide per-cell stripe scores (*right*), colored by annotated cell type. Adjusted Rand index (ARI) values are shown for each embedding. For genome-wide profiling, each stripe was represented by one score per cell using the distal 50% of the stripe body relative to matched flanking regions. All panels use cells from the co-assayed HiRES dataset [24]; cell-type, major-cell-population, cell-cycle and developmental-stage annotations follow the original study.

The matched scRNA-seq profiles in HiRES further allowed us to examine whether genome-wide stripe-score profiles could be integrated with transcriptomic information from the same cells. Joint embedding of per-cell stripe scores and matched scRNA-seq showed broadly concordant major-cell-population structure with the RNA-only embedding (Fig. 5c). Together, these analyses show that genome-wide per-cell stripe scores capture multiple axes of biological variation and can be combined with matched gene-expression profiles. The resulting cell-matched structural and transcriptional measurements provide a basis for subsequent analysis of stripe–gene-expression associations across individual cells.

### Genome-wide integration of per-cell stripe scores with matched scRNA-seq identifies cell-type-specific gene–stripe relationships

The scMicro-C study resolved heterogeneous promoter–enhancer stripe configurations at the *EBF1* locus and other loci across individual cells and proposed that such variation may contribute to transcriptional bursting [21]. However, matched scRNA-seq profiles were unavailable from the same cells in that dataset, precluding direct examination of stripe–expression relationships at the single-cell level. Co-assayed scHi-C and scRNA-seq datasets provide an opportunity to examine these relationships directly in the same cells, although their chromatin-contact maps can be substantially sparser than those generated by scMicro-C. For example, cells in the HiRES dataset contained an average of approximately 160,000 contacts within 2 Mb per cell, compared with approximately 555,000 in the scMicro-C dataset. Building on the locus-level and genome-wide analyses above, we therefore examined relationships between per-cell stripe scores and matched gene expression across individual cells at genome-wide scale. As a proof-of-concept, we used the HiRES dataset [24], with each stripe represented by one score per cell using the genome-wide profiling mode.

We first examined relationships between stripe-anchor proximity and matched gene expression. A gene was considered proximal when its TSS lay within the directional span of a stripe and within 50 kb of the anchor. In blood cells, genes with detectable expression were more frequently proximal to stripe anchors than genes with undetectable expression (Fig. 6a), with a similar pattern observed across cell types (Supplementary Fig. S14a). Consistent with this pattern, proximal genes showed higher expression than non-proximal genes among the union of the top 100 up-regulated genes from each cell type (Fig. 6b, Supplementary Fig. S14b). This expression contrast remained qualitatively stable under alternative proximal definitions (Supplementary Fig. S14c). Extending beyond the fixed proximal definition (directional, 50 kb), the distances from the same up-regulated genes to the nearest stripe anchor were shifted toward shorter values relative to randomly selected genes (Fig. 6c). We next asked whether gene expression was also related to stripe-score magnitude. In blood cells, average gene expression showed a modest positive association with the average per-cell stripe score of the nearest proximal stripe among proximal up-regulated genes (Fig. 6d). Together, these analyses support genome-wide associations between stripe features and matched gene expression.

**Figure 6:**
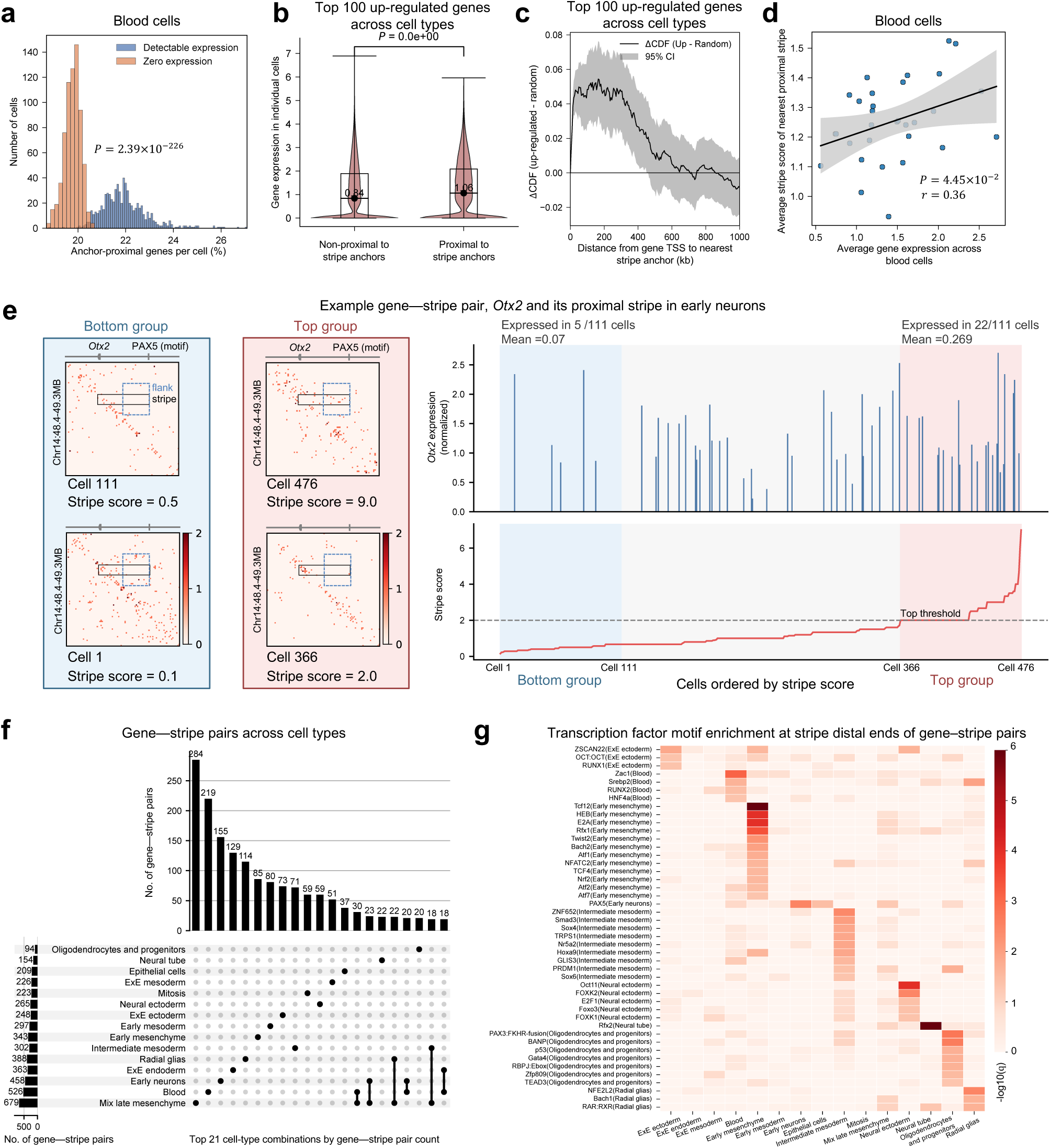
Genome-wide per-cell stripe scores are associated with matched gene expression and identify cell-type-specific gene–stripe relationships. **a,** In blood cells, genes with detectable expression showed a higher per-cell proportion of anchor-proximal genes than genes with zero expression. **b,** Across cell types, the union of the top 100 up-regulated genes from each cell type showed higher expression when genes were proximal rather than non-proximal to stripe anchors. **c,** Difference between the cumulative distribution functions (ΔCDF, up-regulated minus random) of distances from gene transcription start sites (TSSs) to the nearest stripe anchor for the same up-regulated genes and randomly selected genes. Positive ΔCDF values indicate enrichment of shorter distances among the up-regulated genes; shading denotes the 95% confidence interval. **d,** In blood cells, association between average gene expression and the average per-cell stripe score of the nearest proximal stripe among proximal up-regulated genes. Each point represents one gene. **e,** Representative gene–stripe pair comprising *Otx2* and its proximal stripe in early neurons. Left, representative raw scHi-C contact maps from cells in the bottom and top stripe-score groups, with the stripe, flanking region and distal PAX5 motif indicated. Right, normalized *Otx2* expression and per-cell stripe scores across cells ordered by stripe score. Shaded regions mark the size-matched bottom and top groups, and the dashed line marks the stripe-score threshold of 2 used to define the top group. **f**, UpSet plot of gene–stripe pairs across cell types. Vertical bars show gene–stripe pair counts for the leading 21 cell-type combinations, and the dot matrix indicates the corresponding cell types. **g**, Transcription factor motif enrichment at stripe distal ends of gene–stripe pairs across cell types. Heatmap values show *−* log_10_(*q*); only significantly enriched motifs are shown. A gene was considered proximal to a stripe anchor when its TSS lay within the directional span of the stripe and within 50 kb of the anchor. For the gene–stripe pair analyses in **e–g**, cells were ordered by per-cell stripe score. The top group comprised cells with stripe score *≥* 2, subject to the group-size criteria described in Methods, and the bottom group contained an equal number of cells with the lowest stripe scores. Candidate pairs were ranked by fold change in average gene expression between the top and bottom groups, and the top 25% within each cell type were retained as final gene–stripe pairs. All analyses used matched scHi-C and scRNA-seq profiles from the HiRES dataset [24]. Stripes were identified at 10 kb resolution from aggregated scHi-C maps for each cell type, and per-cell stripe scores were computed directly from the corresponding raw single-cell contact maps using the genome-wide profiling mode.

To further examine relationships between per-cell stripe-score variation and gene expression across individual cells, we focused on positively associated gene–stripe relationships within each cell type. A gene–stripe pair was defined when a gene was proximal to a stripe anchor and showed higher expression in the top group than in the size-matched bottom group after cells were ordered by per-cell stripe score. As a representative example, *Otx2* and its proximal stripe formed a gene–stripe pair in early neurons. A total of 111 cells had a per-cell stripe score ≥ 2 for this stripe and were assigned to the top group. Among these cells, *Otx2* transcripts were detected in 22 cells, with a mean expression of 0.269, whereas transcripts were detected in only 5 of the 111 cells in the size-matched bottom group, with a mean expression of 0.07, corresponding to an approximately 3.84-fold difference in average expression (Fig. 6e). Across cell types, 94–679 gene–stripe pairs were identified per cell type. The identified gene–stripe pairs were broadly robust to the stripe-score threshold used to define the top group, with similar pair counts, substantial overlap with the default threshold of 2.0, and consistently positive expression fold changes across thresholds of 1.5, 2.0 and 2.5 (Supplementary Fig. S14d). Consistent with the positive-association criterion, fold changes in average expression between the top and bottom groups across the final gene–stripe pairs ranged from 1.16 to 36.59, with a median of 1.46.

We next examined the cell-type specificity of the identified gene–stripe pairs. UpSet analysis showed that the leading gene–stripe pair combinations were predominantly restricted to a single cell type rather than shared across cell types (Fig. 6f). Across cell types, the proportion of gene–stripe pairs restricted to a single cell type ranged from 14.3% to 41.8%, with a median of 26.5%, approximately fivefold higher than the median of 5.7% for stripes (range, 2.5–12.7%; Fig. 6f; Supplementary Fig. S15). Thus, the identified gene–stripe pairs showed substantially greater cell-type specificity than the stripes themselves. Motif-enrichment analysis at stripe distal ends of gene–stripe pairs identified 1–12 significantly enriched transcription factor (TF) motifs per cell type, with 40 of 45 enriched motifs detected in only one cell type (Fig. 6g). Examples included FoxO3 in neural ectoderm [43], PAX5 in early neurons [44], Tcf12 in early mesenchyme [45], p53 in oligodendrocytes and progenitors [46], and RAR:RXR in radial glias [47] (Fig. 6g). Motif enrichment at stripe anchors showed a similar pattern across cell types (Supplementary Fig. S16).

Together, these analyses show that genome-wide per-cell stripe scores can be integrated with matched scRNA-seq profiles to examine stripe–expression relationships across individual cells and to identify positively associated gene–stripe pairs that show pronounced cell-type specificity and distinct transcription factor motif enrichment across cell types. Because these associations are derived from static single-cell profiles, they do not establish temporal order or causal directionality.

## Discussion

Chromatin stripe analysis in scHi-C is challenging because resolving stripes requires regulatory-scale contact maps, whereas individual-cell maps are extremely sparse at these resolutions. scStripe addresses this challenge with an imputation-free two-stage strategy that detects stripes from sparse cell-type–aggregated scHi-C maps and directly quantifies the resulting aggregate-defined stripes in raw individual-cell maps using per-cell stripe scores. Across diverse datasets, scStripe achieved the highest F1 scores in three complementary benchmark settings, showed strong aggregate stripe enrichment, and recapitulated characteristic structural and regulatory features of stripes reported in bulk data. At the per-cell level, stripe scores resolved recurrent configurations of the *EBF1*-anchored stripe, captured genome-wide variation associated with cell type, cell-cycle phase, and developmental stage, and enabled analysis of stripe–expression relationships with pronounced cell-type specificity using matched scRNA-seq. Together, scStripe links aggregate-level stripe detection to direct per-cell analysis at regulatory resolution without requiring single-cell enhancement or 3D reconstruction.

For aggregate-level stripe detection, scStripe separates candidate identification from statistical evaluation. Random-matrix-theory–guided spectral selection and change-point detection identify candidate endpoints from sparse aggregated scHi-C maps, after which complementary tests evaluate candidate stripes for enrichment relative to matched flanking backgrounds and for sustained enrichment toward their distal ends. This separation allows stripe support to be assessed against complementary matched backgrounds rather than through a single local-background comparison that may be sensitive to sparsity in aggregated scHi-C maps. At the per-cell level, scStripe addresses sparsity by directly quantifying aggregate-defined stripes rather than reconstructing dense contact maps for individual cells. The per-cell stripe score pools contacts across the stripe and matched flanks and quantifies their relative enrichment, thereby aggregating information over many bin pairs while normalizing against matched local backgrounds within the same cell. Because the score is defined as a ratio of pooled stripe and matched-flank contacts, it is expected to be relatively robust to differences in sequencing depth across cells. Downsampling and held-out analyses supported this expectation, showing progressively greater agreement with full-depth stripe-score profiles as sequencing depth increased and limited sensitivity to self-inclusion. Together, these complementary designs enable regulatory-scale stripe detection from sparse aggregated scHi-C maps and direct per-cell quantification from extremely sparse individual-cell maps.

Per-cell stripe scores transform aggregate-defined stripes into quantitative features that can be compared across individual cells. At the *EBF1* locus, within-stripe tiling revealed recurrent configurations that differed primarily in distal reach, resolving heterogeneity within an aggregate-defined stripe that is obscured by aggregation. Genome-wide stripe-score profiles further captured variation associated with cell type, cell-cycle phase, and developmental stage, indicating that stripe-associated contact enrichment varies along multiple biological axes. Using matched scRNA-seq from the same cells, per-cell stripe scores further enabled genome-wide analysis of stripe–expression relationships, identifying positively associated gene–stripe pairs with pronounced cell-type specificity. Together, these analyses show that per-cell stripe scores support both locus-specific and genome-wide characterization of stripe variation and its integration with other single-cell measurements.

Several limitations should be considered when applying scStripe. First, aggregate-level stripe detection requires predefined cell-type labels and depends on the contact coverage available within each aggregated cell population, as reflected by the positive association between stripe counts and sequencing depth (Fig. 2d). Stripes confined to rare cell states or subtypes within a broader cell population could be missed if their signals are diluted below the level detectable in the aggregated map. Second, per-cell stripe scoring is designed to quantify aggregate-defined stripes and does not perform de novo stripe detection or stripe extent estimation in individual cells. Third, stripe–expression relationships identified from matched scRNA-seq are based on static measurements from the same cells and therefore do not establish temporal order or causal directionality. Higher-coverage single-cell chromatin-contact datasets and finer cell-state annotations may enable stripe detection in smaller cell populations and broaden the range of stripes that can be examined at the per-cell level. Time-resolved, imaging, and perturbation studies will be needed to determine the temporal and causal relationships between stripe variation and gene regulation. Within these limitations, scStripe provides a practical, scalable framework that connects regulatory-scale stripe detection in sparse aggregated scHi-C maps with quantitative analysis across individual cells, without requiring single-cell enhancement or 3D reconstruction.

## Methods

### Data preprocessing and normalization

Raw paired-contact files for the single-cell chromatin conformation datasets were obtained from the corresponding public repositories and converted to Cooler format using `cooler cload pairs` [48]. For each annotated cell type, contacts from individual cells were aggregated to generate cell-type–aggregated contact maps. Aggregated maps were normalized using iterative correction and eigenvector decomposition (ICE) as implemented in `cooler balance` [49]. For the Tan et al. dataset [27], ICE normalization did not converge at 10 kb resolution, and Knight–Ruiz (KR) normalization was therefore used instead [50]. The lineage-related NPC bulk Hi-C dataset used for benchmarking was processed using the same workflow and normalized by ICE.

Individual-cell contact maps were generated at 10 kb resolution and analyzed directly using raw, unbalanced contact counts without imputation or smoothing. For the scMicro-C dataset [21], cells with fewer than 200,000 contacts were excluded, leaving 758 cells for analysis; contacts from these cells were also aggregated to generate the corresponding cell-type–aggregated scMicro-C map. Unless otherwise specified, analyses were performed at 10 kb resolution.

### Nomenclature of 5′ - and 3′ -stripes

Chromatin stripes can extend from an anchor toward either lower or higher genomic coordinates. Following the convention used in bulk Hi-C studies [2], stripe orientation is defined with respect to the upper triangle of the contact matrix. A stripe anchored at a genomic locus and extending toward lower genomic coordinates appears as a vertical feature and is defined as a 5′-stripe, whereas a stripe extending toward higher genomic coordinates appears as a horizontal feature and is defined as a 3′-stripe. These labels refer to stripe orientation in the contact map rather than to the transcriptional orientation of nearby genes.

### scStripe

scStripe is an imputation-free two-stage statistical framework for detecting and quantifying chromatin stripes from scHi-C contact maps. In stage 1, scStripe detects stripes from normalized cell-type–aggregated scHi-C maps by applying random-matrix-theory–guided spectral selection to retain informative eigenvectors, change-point detection to identify candidate stripe endpoints, and complementary statistical tests with fold-change criteria to evaluate candidate stripes against matched backgrounds. In stage 2, scStripe quantifies the resulting aggregate-defined stripes directly in raw individual-cell contact maps using per-cell stripe scores, yielding a stripe-by-cell matrix in which each entry measures relative stripe enrichment in one cell. The two stages are performed without single-cell enhancement or 3D reconstruction.

#### Input and output

scStripe accepts either contact matrices or Cooler files (`.cool`) as input. Stage 1 outputs a set of aggregate-defined stripes, each reported by its genomic coordinates together with stripe length and width. Final stripe calls are saved in BEDPE format for downstream analysis and visualization.

Stage 2 outputs a per-cell stripe-score matrix. In genome-wide profiling mode, rows correspond to aggregate-defined stripes and columns to individual cells, with each entry representing the per-cell stripe score for one stripe in one cell. By default, the score is computed from the distal 50% of the stripe body. For within-stripe tiling, a selected stripe can instead be partitioned into multiple non-overlapping subregions, yielding a subregion-by-cell matrix that resolves variation within the stripe.

#### Spectral decomposition and RMT-guided selection of informative eigenvectors

Cell-type–aggregated scHi-C contact maps were partitioned along the main diagonal into overlapping 200*×*200-bin submatrices, corresponding to 2 Mb*×*2 Mb at 10 kb resolution, with a step size of 50 bins. For each submatrix, the contact matrix *M* was eigendecomposed as

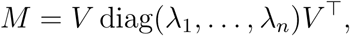

where *λ_i_*and *v_i_*denote the *i*-th eigenvalue and corresponding eigenvector, respectively.

Motivated by random matrix theory, which distinguishes a limited number of structured spectral modes from a bulk of weaker modes [51], scStripe retains leading eigenvectors according to the empirical structure of the eigenvalue spectrum rather than applying a model-specific theoretical eigenvalue cutoff. Eigenvalues are ordered by decreasing absolute magnitude,

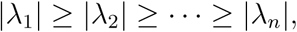

and the number of retained eigenvectors *r* is determined by the first local plateau satisfying

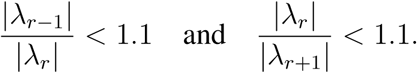

The retained eigenvectors summarize dominant coherent contact patterns in the contact submatrix and may capture stripe-associated signals as well as other chromatin-organization patterns; they are therefore used as low-dimensional inputs for change-point detection rather than interpreted as stripe-specific components.

Each retained eigenvector is analyzed independently using the Pruned Exact Linear Time (PELT) change-point algorithm implemented in `ruptures` with `model=“rbf”` [52, 53]. PELT identifies positions at which the eigenvector profile changes along the genomic axis by minimizing a penalized segmentation cost,

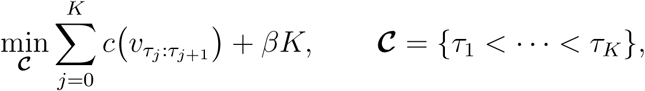

where *β* controls the number of detected change-points. A penalty of 0.1 was used by default. The `jump` parameter was set to 2, such that candidate change-points were evaluated on a two-bin grid along the genomic axis. Change-points identified across the retained eigenvectors were collected and ordered to obtain the candidate stripe endpoints used in the subsequent stripe-construction step (Fig. 1a).

#### Candidate stripe construction

The ordered change-points identified from the retained eigenvectors are used to define candidate stripe geometries. Let *b*_1_ *≤ b*_2_ *≤· · · ≤ b_m_* denote the ordered candidate endpoints within a contact-map submatrix. Each adjacent pair (*b_i_, b_i_*_+1_) defines a candidate width interval near the diagonal, and scStripe then searches unidirectionally from this interval for a distal endpoint that determines stripe length.

To reduce the influence of distance-dependent contact decay, each submatrix is transformed to observed/expected (O/E) values before distal-endpoint selection. For genomic offset *d* = *j−i*, the expected contact value is calculated as the mean contact intensity along the corresponding diagonal,

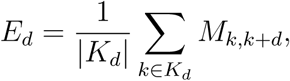

where *K_d_*contains all valid bin pairs at offset *d*, and the O/E value is given by

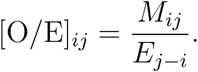

For each candidate width interval, scStripe determines stripe orientation and searches along the corresponding genomic direction for a distal endpoint. For every possible extension, a candidate-generation fold change is calculated as

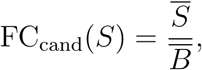

where *S̅* is the mean O/E value within the candidate stripe and *B̅* is the mean O/E value of the corresponding 200*×*200-bin submatrix. The submatrix mean is used at this stage as an efficient background reference for distal-endpoint localization. The distal endpoint is selected as the first prominent local maximum of FC_cand_ that exceeds the predefined threshold, corresponding to the point beyond which extending the candidate no longer increases its relative enrichment.

Adjacent or overlapping candidate stripes in the contact map are subsequently consolidated. Distal endpoints are considered concordant when they differ by no more than half of the minimum stripe length, corresponding to 10 bins under the default minimum length of 20 bins. Candidates with contiguous width intervals and concordant distal endpoints are merged, whereas candidates separated by more than the maximum stripe width are retained separately. Stripe width is further refined using local contact gradients under a bounded expansion constraint. Candidates with width greater than 8 bins or length shorter than 20 bins are discarded by default.

#### Statistical testing of candidate stripes

Candidate stripes retained after the construction and refinement steps are evaluated using three one-sided tests against matched background regions. For clarity, the procedure is described below for a 3′-stripe in the lower triangle of the contact submatrix; the corresponding regions are mirrored for 5′-stripes.

For a candidate stripe *S*, two flanking regions, *L* and *R*, are defined on either side of the stripe and matched to *S* in size, shape, anchor position and distance-to-diagonal profile. These two comparisons test whether contact enrichment within the stripe exceeds the local backgrounds on both sides. A third comparison evaluates the distal termination of the candidate stripe. The terminal segment of the stripe adjacent to the detected distal endpoint is denoted by *S*′, and a matched background region of equal size immediately beyond the distal endpoint is denoted by *D*. The *S*′–*D* comparison tests whether enrichment is retained up to the detected distal endpoint relative to the region beyond it.

For the two flank comparisons, consider a 3*^→^*-stripe in the lower triangle, where the stripe is vertically oriented. At each row corresponding to one position along the stripe length, the O/E contact values across the columns spanning the stripe width are averaged to obtain one stripe value. The same row-wise averaging is applied to the width-matched left and right backgrounds, *L* and *R*. Repeating this operation over all rows along the stripe yields three equal-length one-dimensional profiles, with each element of the three profiles corresponding to the same position along the stripe and therefore forming a matched observation for the paired tests. The resulting stripe and background values are log-transformed, and one-sided paired *t*-tests are performed for

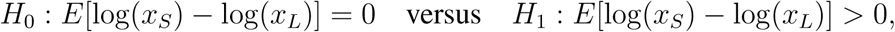

And

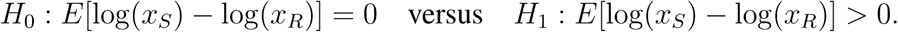

For the distal-end comparison, values within *S′* and *D* are averaged across the stripe width to generate matched profiles along the stripe direction, followed by the analogous one-sided paired *t*-test

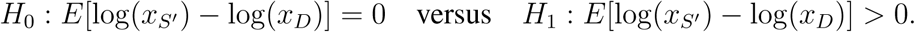

The resulting *P* values are adjusted across candidate stripes using the Benjamini–Hochberg procedure. Candidates are required to satisfy an FDR-adjusted *P* -value threshold of 0.001 for both flank comparisons and 0.05 for the distal-end comparison.

In addition to statistical significance, the two flank comparisons are required to satisfy minimum fold-change criteria. Fold changes are calculated from mean O/E contact values as

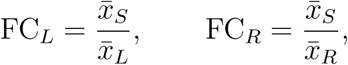

where *x̅_S_*, *x̅_L_* and *x̅_R_* denote the mean O/E values within the stripe and its two matched flanking regions. A minimum fold change of 1.1 is required for both comparisons. Candidate stripes satisfying all three significance criteria and both flank fold-change criteria are retained for subsequent de-duplication.

#### Post-processing and de-duplication

Candidate stripes satisfying the statistical and fold-change criteria are subsequently de-duplicated to remove redundant calls arising from overlapping contact-map patches or closely related candidate geometries. The procedure was adapted from Stripenn [31]. Stripes are first grouped by chromosome and compared within the local genomic windows used by the pipeline.

For each pair of stripes, overlap is quantified separately along the stripe-width and stripe-length dimensions. Two stripes are considered redundant when the overlap proportion exceeds 0.2 along both dimensions. For each redundant pair, the stripe with the smaller length-to-width ratio is removed. The remaining nonredundant stripes constitute the final stripe set for the corresponding cell-type–aggregated contact map and are saved in BEDPE format for downstream analyses.

#### Per-cell stripe scoring

In stage 2, scStripe quantifies aggregate-defined stripes directly in raw individual-cell contact maps without imputation or smoothing. For an aggregate-defined stripe *S* and cell *c*, the corresponding raw contact map is binarized at the analysis resolution, and *n_c_*(*S*) denotes the number of observed bin-pair contacts within *S*. Let *L* and *R* denote the two flanking regions matched to *S* in size and shape, with *n_c_*(*L*) and *n_c_*(*R*) denoting their corresponding contact counts. The per-cell stripe score is defined as

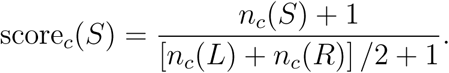

The score therefore compares pooled contacts within the stripe with the average pooled contact count in its two matched flanks.

Raw individual-cell scHi-C contact maps are extremely sparse at 10 kb resolution, such that contact observations at individual bin pairs are frequently absent. The per-cell stripe score is designed for this sparse setting by pooling contacts across multiple bin pairs within the stripe and its matched flanks, thereby aggregating information over the full scoring region rather than relying on individual contact observations. The score then expresses the pooled stripe signal relative to the matched local background within the same cell. Because the stripe and flanking regions are observed from the same cell, variation in overall sequencing depth is expected to affect the numerator and denominator in the same direction, reducing the influence of cell-to-cell differences in contact coverage. The +1 pseudocount prevents undefined ratios when flank contacts are absent and limits instability when pooled counts are very small. This expected robustness to extreme sparsity and sequencing depth was evaluated empirically by the contact-downsampling analyses described in Supplementary Methods.

The per-cell stripe score is distinct from the fold-change quantities used during stage 1. During candidate construction, candidate-generation fold change is calculated relative to the corresponding contact-map patch and is used to localize the distal endpoint. During final statistical filtering, fold changes are calculated relative to the two matched flanking backgrounds. By contrast, the stage-2 per-cell stripe score is calculated directly from raw individual-cell contact maps and measures relative stripe enrichment using pooled contact counts.

For genome-wide profiling, one per-cell stripe score is calculated for every aggregate-defined stripe in every cell. By default, scoring is restricted to the distal 50% of the stripe body, together with the corresponding portions of its matched flanks. This restriction reduces the influence of strong near-diagonal contact enrichment, which can be shared by the stripe and its flanking regions and thereby reduce contrast between them. The resulting stripe-by-cell matrix provides one quantitative feature for each stripe in each cell and is used for downstream analyses including marker-stripe identification, cell embedding and integration with matched scRNA-seq profiles.

For within-stripe tiling, a selected stripe is partitioned along its anchor-to-distal axis into multiple non-overlapping subregions. The same per-cell stripe-score definition is then applied separately to each subregion and its corresponding matched flanks, yielding a subregion-by-cell matrix. This mode retains positional variation along the stripe that would be summarized by a single stripe-level score in genome-wide profiling.

### Stripe-calling methods used for benchmarking

We compared scStripe with six alternative stripe callers: Stripenn [31], StripeDiff [32], JOnTADS [33], StripePy [34], StripeCaller and Zebra [2]. For the large-scale cross-cell-type benchmark, scStripe, Stripenn, StripeDiff, JOnTADS, StripePy and StripeCaller were applied at 10 kb resolution to the same normalized cell-type–aggregated scHi-C contact maps. Default settings or configurations recommended by the corresponding publications were used unless otherwise specified. Zebra requires a distinct preprocessing workflow and was therefore included only in focused comparisons. Following stripe detection, calls were restricted to stripes with length ≥ 20 bins and width ≤ 8 bins, corresponding to lengths of at least 200 kb and widths of at most 80 kb at 10 kb resolution.

StripeDiff identifies stripe anchors using an edging signal that contrasts contact intensities across candidate stripe boundaries and estimates stripe extent by applying change-point detection to the resulting peak-intensity profiles [32]. We used the recommended configuration, including the default line-scan window size of 300 bins.

Stripenn uses image-processing procedures including contrast adjustment and Canny edge detection to identify linear contact-enrichment features [31]. Candidate stripes are evaluated using a median *P* value that measures contrast with neighboring regions and a stripiness score that characterizes stripe continuity and contrast. We applied the default filtering scheme with median *P <* 0.1.

JOnTADS jointly identifies TADs and stripes by scanning contact maps for linear enrichment patterns and refining stripe extent using a cumulative enrichment statistic. Following the original implementation, detected stripes were filtered using the Stripenn-derived significance measure with *P <* 0.1.

StripePy identifies candidate stripes from one-dimensional marginal contact profiles using topological-persistence-based detection of local maxima and subsequently estimates stripe width and length from the surrounding contact pattern [34]. StripePy was run using the recommended default settings.

StripeCaller is an independently implemented stripe caller based on the Zebra procedure and was included in the large-scale cross-cell-type benchmark using its default settings [31]. For Zebra, the source code was obtained directly from the corresponding author and run using the required preprocessing workflow. The original Zebra study additionally applied manual post-processing to the detected stripe calls. Because such manual curation is infeasible for systematic benchmarking across many cell types and may introduce subjective bias, we used the stripe calls produced directly by the Zebra implementation without additional manual post-processing. Zebra was therefore evaluated only in focused comparisons rather than in the full cross-cell-type benchmark.

### Benchmarking analysis

#### Cross-cell-type benchmarking design

Stripe detection was evaluated using cell-type–aggregated contact maps from 94 annotated cell types across nine single-cell chromatin-conformation datasets. For each cell type, all methods included in the large-scale benchmark were applied to the same normalized aggregated contact map. The number of cell types in which each method detected at least one stripe was first recorded across all 94 cell types.

For direct cross-method comparisons of stripe counts, overlap and aggregate enrichment, analyses were restricted to cell types in which all six methods included in the large-scale benchmark, scStripe, Stripenn, StripeDiff, JOnTADS, StripePy and StripeCaller, detected at least one stripe. This criterion yielded 42 cell types across eight datasets. Zebra was not included in this filtering criterion because it was evaluated only in focused comparisons.

#### Common and unique stripes

To quantify overlap between stripe call sets, two stripes were considered overlapping when their genomic regions shared at least one genomic bin on the same chromosome, following the overlap definition used in Stripenn [31]. For the focused comparison at the nine age-associated marker-gene loci in excitatory neurons, stripes detected by the alternative callers were merged into a combined reference set and classified according to whether they overlapped a scStripe call. Genome-wide overlap between scStripe and individual alternative callers was evaluated using the same criterion.

#### Concordance with lineage-related bulk Hi-C

Concordance between stripes detected from the excitatory-neuron aggregated scHi-C map and lineage-related NPC bulk Hi-C was evaluated separately for each method. For each stripe caller, the corresponding reference set was generated by applying the same method to the NPC bulk Hi-C map, thereby avoiding comparison of one method against stripe calls produced by another method.

For scStripe and the alternative callers except JOnTADS, stripe sets from the aggregated scHi-C and bulk Hi-C maps were rasterized into binary contact-map matrices at the corresponding resolution. Pixels covered by at least one stripe were assigned a value of 1 and all remaining pixels a value of 0. Pixel-level precision was defined as the fraction of stripe-covered pixels in the aggregated scHi-C map that were also covered by stripes in the bulk Hi-C reference,

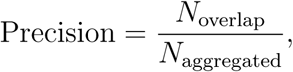

and pixel-level recall as the fraction of stripe-covered pixels in the bulk Hi-C reference that were recovered in the aggregated scHi-C map,

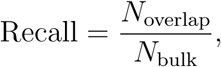

where *N*_overlap_ denotes the number of pixels covered by both stripe sets, and *N*_aggregated_ and *N*_bulk_ denote the numbers of stripe-covered pixels in the aggregated and bulk maps, respectively. The F1 score was calculated as the harmonic mean of precision and recall.

JOnTADS produces stripes with a fixed width of one bin, making pixel-level overlap disproportionately sensitive to small positional differences. For JOnTADS, concordance was therefore evaluated at the stripe level. A stripe detected from the aggregated scHi-C map was considered supported when its rectangular genomic span overlapped a JOnTADS stripe detected from the NPC bulk Hi-C map along both genomic axes. Precision and recall were then defined as the fractions of aggregated-map and bulk-map stripes, respectively, with at least one supported overlap, and the F1 score was calculated from these stripe-level precision and recall values.

#### Simulation benchmark with SimuStripe

Controlled simulations were generated using SimuStripe following the published procedure without modification [32]. SimuStripe generates contact matrices containing predefined embedded stripes, which were used as the reference calls for evaluating stripe detection. We generated 100 simulated contact maps and applied scStripe, StripeDiff and JOnTADS to each simulation. These three methods were included because they can operate directly on the simulated contact matrices; Stripenn, StripePy, StripeCaller and Zebra require Cooler-format or whole-chromosome inputs and were therefore not evaluated in this benchmark. Precision, recall and F1 score were calculated for each simulation against the predefined embedded stripes and summarized across the 100 simulations.

#### Benchmarking against literature-reported stripes

To provide an independent benchmark based on previously reported chromatin stripes, we manually curated 55 stripes from seven published bulk Hi-C studies. A stripe was included when it was explicitly described as a chromatin stripe in the original study, the corresponding contact map was shown in the main text or supplementary material, and its genomic coordinates could be unambiguously determined from the publication. The curated stripe coordinates, corresponding bulk Hi-C datasets, species and source publications are provided in Supplementary Table S1.

The corresponding high-coverage bulk Hi-C contact maps were analyzed at resolutions appropriate to the source datasets rather than at a single fixed resolution. This choice accommodates substantial heterogeneity in both stripe geometry and sequencing depth across the published datasets. The curated stripes had widths of approximately 18–130 kb and lengths of approximately 300 kb–2.72 Mb.

scStripe, StripeDiff and JOnTADS were evaluated on the full set of 55 literature-reported stripes. Four corresponding bulk Hi-C datasets were not available in the input format required by Stripenn, StripePy, StripeCaller and Zebra, leaving 51 literature-reported stripes for the seven-method comparison. For each caller, precision, recall and F1 score were calculated by comparing the detected stripes with the curated literature-reported stripes in their corresponding bulk Hi-C maps.

#### Aggregated stripe analysis

Aggregated stripe analysis (ASA) was used to quantify average contact enrichment around stripes detected by each method. Stripe-centered contact-map regions were extracted and aggregated using `coolpup.py` with the options `--local --rescale --rescale flank 1`. Each stripe-centered region was rescaled to a common coordinate system spanning one stripe length on either side of the stripe, allowing stripes of different genomic lengths to be averaged. ASA was reported only for cell type–method combinations containing at least one detected stripe.

The ASA score was defined as the ratio of the mean normalized contact intensity within the central stripe submatrix to that within the upper-right flanking submatrix,

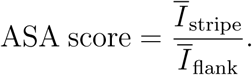

An ASA score greater than 1 therefore indicates higher average contact enrichment within the stripe region than in the matched flanking background. ASA was calculated separately for each method and cell type using the corresponding detected stripe set.

#### Computation time and peak memory

Two benchmarking analyses were performed to evaluate computational requirements. First, runtime and peak memory usage for stripe detection were compared across methods using chromosome 1 from each cell-type–aggregated scHi-C contact map. To ensure comparability, all methods were run using a single CPU thread, with method-specific multithreading disabled when applicable.

Second, the computational requirements of scStripe stage 2 were evaluated independently for genome-wide per-cell stripe scoring across all chromosomes. Cells were processed in batches of 40, and runtime and peak memory were recorded for each cell type. This analysis was intended to assess the scalability of per-cell stripe scoring rather than to provide a cross-method comparison.

Runtime and peak memory usage were measured using `/usr/bin/time -v`. All benchmarks were performed on a Linux server equipped with two Intel Xeon Gold 6240 CPUs (2.60 GHz; 72 threads in total) and 512 GB RAM running CentOS 7.

### Compartment analysis in aggregated scHi-C maps

A/B compartments were identified from cell-type–aggregated scHi-C maps at 50 kb resolution using HOMER [54]. The first principal component (PC1) was oriented using gene transcription start-site annotations such that positive and negative PC1 values corresponded to A and B compartments, respectively. To characterize stripe–compartment relationships, each stripe was classified according to the compartments traversed by its genomic span. Stripes lying entirely within A compartments were classified as only A, those lying entirely within B compartments as only B, and those spanning both compartment types as mixed. For comparison across callers while limiting differences arising from call-set size, up to 200 stripes per method and cell type were retained. Stripes were ranked by Stripenn-derived *P* values, and all stripes were retained when fewer than 200 were detected.

To assess whether stripes in different compartment classes retained stripe-associated contact enrichment, aggregated stripe analysis (ASA) was performed separately for only A and only B stripes. TADs classified entirely within A or B compartments were analyzed in parallel as a structural reference. Compartment-stratified ASA was restricted to stripe or TAD sets meeting the minimum feature-count criterion used for ASA.

### TAD detection and stripe–TAD analysis

TADs were identified from cell-type–aggregated scHi-C maps at 10 kb resolution using the insulation-score method implemented in `cooltools insulation` with a 2 Mb sliding window [42]. Low-quality bins identified by `cooltools` were excluded. Positions with positive insulation boundary strength were retained as TAD boundaries, and consecutive boundaries were paired to define TAD intervals.

For the Dip-C dataset of Tan et al. [27], ICE normalization did not converge at 10 kb resolution and KR normalization was used instead. Because `cooltools insulation` does not support the KR-normalized maps used here, the 11 Dip-C-derived cell types were excluded from analyses requiring TAD annotations. This left 31 cell types across seven datasets. The same 31-cell-type set was used for the cross-cell-type compartment summaries to maintain direct comparability between the compartment and TAD analyses.

To quantify stripe–TAD relationships, the genomic span of each detected stripe from its anchor to distal end was intersected with the TAD intervals called from the same aggregated scHi-C map. The number of TADs overlapped by each stripe was recorded as the number of TADs spanned by that stripe. Unlike the compartment analysis, which used up to 200 stripes per method and cell type, the stripe–TAD analysis included all stripes detected by each method.

### CTCF and H3K4me3 enrichment analyses

CTCF and H3K4me3 ChIP–seq data from lineage-related NPCs were used to characterize regulatory features associated with stripes detected from the excitatory-neuron aggregated scHi-C map. Two complementary analyses were performed.

First, TADs were classified separately for the stripe calls from each method as stripy when they overlapped at least one detected stripe and as non-stripy otherwise, following the definition used in previous work [31]. CTCF and H3K4me3 enrichment around TAD boundaries was then summarized separately for stripy and non-stripy TADs as the average ChIP–seq peak count per 10 kb bin centered on the corresponding TAD boundaries.

Second, CTCF and H3K4me3 ChIP–seq signals were aligned directly to stripe coordinates for aggregate visualization. Signal values were extracted from bigWig tracks using custom Python code. Before alignment, stripe coordinates were oriented consistently from the anchor toward the distal end. For each stripe, the interval between the anchor and distal end was rescaled to 100 bins using length-weighted linear interpolation of the stepwise bigWig signal. Flanking regions of 10 kb on either side were each rescaled to 50 bins using the same procedure. The two flanks and rescaled stripe body were concatenated to yield a 200-bin signal profile for each stripe, with missing values set to zero. Stripe profiles were sorted by total signal intensity and visualized as heatmaps. Line plots above the heatmaps show, at each relative position, the proportion of stripes with signal intensity above the heatmap-wide median.

### Per-cell analysis of the *EBF1*-anchored stripe

The *EBF1*-anchored stripe was analyzed using the scMicro-C dataset of Wu et al. [21]. After the cell filtering described above, 758 GM12878 cells were retained and analyzed at 5 kb resolution. scStripe was first applied to the aggregated scMicro-C contact map to identify the approximately 500 kb promoter-anchored stripe at the *EBF1* locus. To resolve variation along the stripe, the stripe body was partitioned into six consecutive non-overlapping tiled subregions guided by the positions of the ABC-predicted enhancers reported in the original study. The subregions were ordered from the distal end toward the stripe anchor, and a per-cell stripe score was calculated separately for each subregion using its corresponding matched flanks, yielding a six-dimensional stripe-score profile for each cell.

For descriptive analyses of subregion retention, a tiled subregion was considered retained in a cell when its per-cell stripe score was at least 1.5. Robustness to this threshold was evaluated by repeating the analysis across thresholds from 1.1 to 2.0. The six tiled stripe scores were also used for hierarchical clustering of the 758 cells, and the resulting dendrogram was partitioned into seven groups for comparison of recurrent stripe-score configurations.

For each group, a group-aggregated 5 kb contact map was generated by summing raw contacts across cells in the group and dividing the resulting contact counts by the number of cells. These maps were used to compare the contact patterns corresponding to the score-defined groups. ABC-predicted enhancer annotations and scMicro-C data were obtained from Wu et al. [21]; CTCF, RAD21 and H3K27ac tracks were obtained from 4DN.

### Genome-wide analysis of per-cell stripe-score profiles

Genome-wide per-cell stripe-score profiles were analyzed using the HiRES dataset [24]. For each cell, one score was calculated for every aggregate-defined stripe using the distal 50% of the stripe body, as described above. Stripe-score values equal to 1, corresponding to no relative enrichment between the stripe and its matched flanks, were set to 0 before downstream analysis, whereas values below or above 1 were retained. After this transformation, cells with fewer than 3,000 non-zero stripe-score entries were removed, and stripe features with non-zero values in fewer than 300 cells were excluded.

The resulting stripe-by-cell matrix was normalized by total stripe-score signal per cell, log-transformed and scaled, followed by principal-component analysis. The first 20 principal components were used to construct a *k*-nearest-neighbour graph with *k* = 35. Louvain clustering was performed at a resolution of 1, and cells were visualized using UMAP. The resulting embedding was annotated using the cell-type, major-cell-population, cell-cycle-phase and developmental-stage annotations provided in the original HiRES study.

Cell-type marker stripes were identified from the processed stripe-score matrix using group-wise differential analysis implemented with `scanpy.tl.rank genes groups` using a *t*-test. *P* values were adjusted using the Benjamini–Hochberg procedure, and stripes with adjusted *P <* 0.05 and positive log-fold changes were retained as marker stripes. For heatmap visualization, per-cell stripe scores for the selected marker stripes were standardized across cells.

To assess reproducibility in an independent dataset, the same genome-wide stripe-scoring and downstream analysis workflow was applied to the dscHi-C dataset [22]. Marker-stripe identification and stripe-score-based UMAP embedding were performed using the annotated brain cell types from the original study.

### Joint embedding of per-cell stripe scores and matched scRNA-seq

Per-cell stripe scores and matched scRNA-seq profiles from the HiRES dataset were integrated at the individual-cell level. For the stripe modality, stripe-score values equal to 1 were set to 0 before downstream analysis, as described above. For each modality, cells with fewer than 2,500 non-zero entries were removed, and features with non-zero values in fewer than 300 cells were excluded. The resulting matrices were normalized by the total signal per cell, log-transformed, and subjected to highly variable feature selection, retaining up to 5,000 features per modality. Each modality was then scaled and analyzed by principal-component analysis.

For the RNA-only analysis, the first 35 principal components were used as the low-dimensional representation. For the joint analysis, only cells passing the filtering criteria in both modalities were retained. The first 15 principal components from the stripe-score modality and the first 20 principal components from the RNA modality were concatenated to generate a 35-dimensional joint representation for each cell. A *k*-nearest-neighbour graph with *k* = 35 was constructed from either the RNA-only or joint representation, followed by Louvain clustering at a resolution of 1.5 and UMAP visualization. All preprocessing, dimensionality reduction, graph construction, clustering and visualization were performed using Scanpy [55].

### Gene–stripe association and motif-enrichment analyses

Matched scRNA-seq profiles from the HiRES dataset were analyzed together with per-cell stripe scores from the same cells. Within each cell type, genes expressed in fewer than 10% of cells were excluded. RNA counts were normalized to a total of 10,000 counts per cell and log-transformed. Gene transcription start sites (TSSs) were obtained from the corresponding genome annotation.

A gene was considered proximal to a stripe when its TSS lay within the directional span of the stripe and within 50 kb of the stripe anchor. This directional definition was applied according to stripe orientation, such that only genes located toward the stripe interior from the anchor were considered proximal. Unless otherwise specified, this 50 kb directional criterion was used throughout the gene–stripe analyses.

To examine the relationship between stripe-anchor proximity and gene expression, genes with detectable and undetectable expression were compared according to the proportion that were proximal to stripe anchors. We further identified the top 100 up-regulated genes from each cell type and pooled their union across cell types. Expression levels of these genes were compared between those proximal and non-proximal to stripe anchors. Robustness to the proximity definition was assessed using alternative anchor-distance thresholds. For the same set of up-regulated genes, distances from each gene TSS to the nearest stripe anchor were compared with those of randomly selected genes. In addition, among proximal up-regulated genes, average gene expression across cells was compared with the average per-cell stripe score of the nearest proximal stripe.

For gene–stripe pair analysis, all proximal gene–stripe combinations were evaluated within each cell type. Cells were ordered by the per-cell stripe score of the corresponding stripe. Cells with stripe scores greater than or equal to 2 were assigned to the top group, subject to a maximum group size of 30% of cells in the cell type and a minimum of 10 cells. A size-matched bottom group was defined using the same number of cells with the lowest stripe scores. For each candidate gene–stripe pair, the fold change in average gene expression between the top and bottom groups was calculated. Candidate pairs were ranked by this fold change within each cell type, and the top 25% were retained as gene–stripe pairs. Robustness was assessed by repeating the analysis using stripe-score thresholds of 1.5, 2.0 and 2.5 to define the top group.

For motif-enrichment analysis, gene–stripe pairs within each cell type were used to define foreground genomic regions, whereas the bottom 25% of candidate gene–stripe pairs ranked by expression fold change were used as the background. Anchor and distal regions were analyzed separately. Distal regions were defined as the 20 kb interval extending inward from the distal end along the stripe axis. Anchor and distal regions were partitioned into 200-bp bins, duplicate intervals were removed, and genomic intervals shared between foreground and background sets were excluded from both. Motif enrichment was then performed using HOMER [54].

## Data availability

All datasets used in this study are publicly available. Processed dscHi-C, Dip-C, GAGE-seq, LiMCA, HiRES (scHi-C and scRNA-seq), and scMicro-C data were downloaded from the Gene Expression Omnibus (GEO) under accession codes GSE285812, GSE162511, GSE238001, GSE240128, GSE223917, and GSE281150, respectively. ChAIR data were obtained from the Genome Sequence Archive (GSA) in the National Genomics Data Center under accession number PRJCA024774. Bulk Hi-C and ChIP–seq (CTCF and H3K4me3) data for NPCs were downloaded from GEO under accession code GSE96107. All relevant processed data supporting this study are available upon reasonable request.

## Code availability

The source code for scStripe is available on GitHub at https://github.com/LiLing228/scStripe.

## Acknowledgements

This work was supported by the National Natural Science Foundation of China (grant 12271536 to D.T.). We also acknowledge the High-performance Computing Public Platform (Shenzhen Campus) of Sun Yat-sen University for computational support.

## Author contributions

Conceptualization: D.T.; Methodology: L.L., Z.L., G.P., C.Y. and D.T.; Software: L.L.; Investigation: L.L., Y.L., H.Z., W.G., Z.L., D.L., G.P., C.Y. and D.T.; Visualization: L.L. and Y.L.; Writing – original draft: L.L. and Y.L.; Writing – review & editing: H.Z., C.Y. and D.T.; Supervision: G.P., C.Y. and D.T.; Funding acquisition: D.T.

