## Supplemental File 1 for "Regulatory-scale stripe analysis from single-cell Hi-C with scStripe"

### Supplemental Information

#### A Supplementary Methods

##### Assessment of stripe robustness across filtering thresholds

To evaluate the robustness of scStripe stripe detection to changes in filtering thresholds, all excitatory-neuron cells from the Wu et al. dataset [22] were aggregated to generate a common cell-type–aggregated scHi-C contact map. The complete scStripe workflow was rerun under each parameter setting.

Three filtering parameters were evaluated: the flank fold-change threshold, the flank $P$-value threshold and the distal-end $P$-value threshold, with default values of 1.1, 0.001 and 0.05, respectively. For each analysis, one parameter was varied while the other two were held at their default values.

For the flank fold-change analysis, the threshold was varied among 1.05, 1.10, 1.15 and 1.20. For the flank $P$-value analysis, the threshold was varied among 0.1, 0.05, 0.01, 0.001 and 0.0001. For the distal-end $P$-value analysis, the threshold was varied over the same range from 0.1 to 0.0001.

Within each parameter series, the stripe set obtained under the most stringent threshold was used as the reference set. Specifically, reference stripe sets were defined using a flank fold-change threshold of 1.20, a flank $P$-value threshold of 0.0001 or a distal-end $P$-value threshold of 0.0001, depending on the parameter being evaluated.

For each parameter setting, all detected stripes were recorded. A stripe detected under a more permissive setting was classified as shared with the reference set when its genomic region completely contained at least one reference stripe; otherwise, it was classified as detected only under the more permissive setting. The total number of detected stripes and the number detected only under the more permissive setting were recorded.

Retention of the reference stripe set was quantified as the percentage of reference stripes whose genomic regions were completely contained within at least one stripe detected under the parameter setting being evaluated, calculated as $N_{\text{covered}}/N_{\text{strict}} \times 100\%$, where $N_{\text{strict}}$ is the total number of stripes in the reference set and $N_{\text{covered}}$ is the number recovered under the evaluated parameter setting.

##### Downsampling analysis of stripe detection and per-cell stripe scoring

###### *Single-cell contact downsampling*

To evaluate the robustness of stripe detection and per-cell stripe quantification across sequencing depths, excitatory neurons from the Wu et al. dataset [22] were subjected to independent random contact downsampling. For each cell, raw contacts were sampled without replacement at sampling depths of 0.10, 0.25, 0.50 and 0.75, with five independent replicates generated at each depth. The full-depth condition (sampling depth = 1.00) corresponded to the original raw single-cell contact data.

All excitatory-neuron cells were randomly partitioned into two equal-sized, non-overlapping groups, denoted group A and group B, and this partition was kept fixed throughout the analysis. For each sampling depth and replicate, downsampled contacts were aggregated separately across cells in groups A and B to generate two corresponding aggregated scHi-C contact maps. These maps were used for stripe detection and subsequent analyses of stripe-calling concordance and per-cell stripe-score robustness.

###### *Stripe-calling stability across sampling depths*

Stripe-calling stability across sampling depths was evaluated using the F1 score and pixel-level Jaccard index. For each sampling depth and replicate, aggregated scHi-C contact maps were generated independently from groups A and B, and each evaluated stripe caller was applied separately to the two maps. For each method, concordance between the resulting stripe sets was quantified using the precision, recall and F1 definitions described for concordance with lineage-related bulk Hi-C in the main Methods.

As a complementary overlap measure, each stripe set was rasterized into a binary contact-map matrix, with pixels covered by at least one stripe assigned a value of 1 and all remaining pixels assigned 0. The pixel-level Jaccard index was defined as

$$J = \frac{N_{\text{intersection}}}{N_{\text{union}}},$$

where $N_{\text{intersection}}$ is the number of pixels covered by stripes in both groups and $N_{\text{union}}$ is the number of pixels covered by stripes in either group. The Jaccard index ranges from 0 to 1, with larger values indicating greater concordance between the two stripe sets.

##### *Fixed stripe sets*

Full-depth aggregated scHi-C contact maps were generated separately from the raw contacts of group A and group B. scStripe was applied independently to the two maps, yielding two fixed stripe sets, denoted as $S_A$ and $S_B$, respectively. These stripe sets were identified only once from the full-depth data and were kept fixed across all sampling depths and downsampling replicates in the subsequent per-cell stripe-score analyses.

##### *Definition of held-out and within-group scoring*

For each fixed stripe set, per-cell stripe scores were calculated using cells from both groups. Specifically, $S_A$ was scored in cells from group A and group B, and $S_B$ was likewise scored in cells from both groups.

Within-group scoring refers to quantifying a fixed stripe set in cells from the same group used to identify that stripe set. Thus, $S_A$ was scored in group A cells and $S_B$ in group B cells. Because these cells contributed to the aggregated contact map from which the corresponding stripe set was identified, within-group scoring may potentially be influenced by self-inclusion.

Held-out scoring refers to quantifying a fixed stripe set in cells from the complementary group, whose cells did not contribute to identification of that stripe set. Thus, $S_A$ was scored in group B cells and $S_B$ in group A cells. This design provides a held-out assessment of per-cell stripe-score robustness and allows the potential influence of self-inclusion to be evaluated.

##### *Stability of per-cell stripe scores across sampling depths*

For each cell, a per-cell stripe-score vector was obtained at sampling depth $d$,

$$\mathbf{S}^{(d)} = \left( s_1^{(d)}, s_2^{(d)}, \dots, s_n^{(d)} \right),$$

where $n$ denotes the number of stripes in the corresponding fixed stripe set. This vector was compared with the stripe-score vector obtained for the same cell under the full-depth condition,

$$\mathbf{S}^{(1.0)} = \left( s_1^{(1.0)}, s_2^{(1.0)}, \dots, s_n^{(1.0)} \right).$$

Per-cell stripe-score consistency was quantified using the Spearman correlation

$$\rho_{\text{cell}} = \text{Spearman} \left( \mathbf{S}^{(d)}, \mathbf{S}^{(1.0)} \right).$$

For each sampling depth, $\rho_{\text{cell}}$ was calculated separately under within-group and held-out scoring for each cell and each downsampling replicate. The resulting correlations were then summarized across the five independent replicates.

##### *Cell–cell distance consistency*

To assess whether relative relationships among cells were preserved after downsampling, pairwise cell–cell distances were calculated from per-cell stripe-score profiles. For each sampling depth and fixed stripe set, the stripe-by-cell score matrix was denoted by

$$M^{(d)} \in \mathbb{R}^{n \times c},$$

where $n$ is the number of stripes and $c$ is the number of cells. For cells $i$ and $j$, the Euclidean distance between their stripe-score profiles was calculated as

$$\delta_{ij}^{(d)} = \sqrt{\sum_{k=1}^n \left( s_{ki}^{(d)} - s_{kj}^{(d)} \right)^2},$$

where $s_{ki}^{(d)}$ denotes the score of stripe $k$ in cell $i$ at sampling depth $d$.

This procedure yielded a cell–cell distance matrix for each sampling depth. To quantify preservation of cell–cell relationships, the upper-triangular elements of the distance matrix, excluding the diagonal, were extracted and compared with the corresponding elements from the full-depth distance matrix. Consistency was quantified using the Spearman correlation

$$\rho_{\text{distance}} = \text{Spearman} \left( \mathbf{u}^{(d)}, \mathbf{u}^{(1.0)} \right),$$

where $\mathbf{u}^{(d)}$ and $\mathbf{u}^{(1.0)}$ denote the vectors of pairwise cell–cell distances at sampling depth $d$ and full depth, respectively. The analysis was performed separately under within-group and held-out scoring for each downsampling replicate.

##### *Stripe-score variability ratio*

To assess whether cell-to-cell variability in per-cell stripe scores was preserved after downsampling, the variance of each stripe score across cells was calculated at sampling depth $d$ and at full depth, denoted by $\text{Var}_i^{(d)}$ and $\text{Var}_i^{(1.0)}$ for stripe $i$, respectively.

A variability ratio was then defined as

$$R_i = \frac{\text{Var}_i^{(d)}}{\text{Var}_i^{(1.0)}}.$$

Values of $R_i$ close to 1 indicate that the degree of cell-to-cell stripe-score variability is preserved relative to the full-depth data, whereas values below or above 1 indicate reduced or increased variability after downsampling, respectively. The ratio was calculated separately under within-group and held-out scoring for each downsampling replicate and summarized across the five independent replicates.

##### *Quantification of self-inclusion effects*

For each cell, per-cell stripe-score consistency was calculated separately under held-out and within-group scoring, yielding $\rho_{\text{held-out}}$ and $\rho_{\text{within-group}}$, respectively. The magnitude of the self-inclusion effect was quantified as the absolute difference

$$\Delta\rho = |\rho_{\text{held-out}} - \rho_{\text{within-group}}|.$$

Values of $\Delta\rho$ close to zero indicate similar stripe-score consistency under held-out and within-group scoring and suggest limited sensitivity of this measure to self-inclusion. For each sampling depth, $\Delta\rho$ was calculated for each cell and summarized across the five independent downsampling replicates.

#### B Supplementary Tables

**Table S1: Description of manually curated stripes and source bulk Hi-C datasets from seven publications.** This table lists a manually curated set of chromatin stripes reported in seven published bulk Hi-C studies, together with their genomic coordinates, dataset accessions (GEO/4DN), species, and original references. Stripes were included only when explicitly described in the original study and supported by a corresponding Hi-C heatmap visualization. Full inclusion criteria are detailed in the Methods. This curated collection serves as a reference set for controlled benchmarking and qualitative validation of stripe-detection methods. In total, the table includes 55 stripes drawn from bulk Hi-C datasets spanning human and mouse across seven independent studies. Resolutions are not fixed to 10 kb, but instead follow those used in the original publications. The curated stripes and their corresponding bulk Hi-C maps are visualized in Supplementary Fig. S9. Additional materials, including stripe-centered contact matrices, the resolution used for each stripe, and associated metadata, are available at https://github.com/LiLing228/gold-standard-stripes.

| N | Stripe region | GEO/4DN | Species | Reference |
| --- | --- | --- | --- | --- |
| 1 | Chr15: 25320001–25380000<br>Chr15: 24500001–25380000 | GSE82144 | mouse |  |
| 2 | Chr12: 114400001–114480000<br>Chr12: 114400001–117260000 | GSE82144 | mouse |  |
| 3 | Chr4: 32090001–32120000<br>Chr4: 32090001–32640000 | GSE82144 | mouse |  |
| 4 | Chr4: 32180001–32220000<br>Chr4: 32180001–32640000 | GSE82144 | mouse |  |
| 5 | Chr4: 32280001–32320000<br>Chr4: 32280001–32640000 | GSE82144 | mouse |  |
| 6 | Chr3: 187430001–187470000<br>Chr3: 187430001–188890000 | GSE63525 | human | Vian et al. [2] |
| 7 | Chr9: 30890001–30920000<br>Chr9: 30240001–30920000 | GSE82144 | mouse |  |
| 8 | Chr4: 64140001–64220000<br>Chr4: 63100001–64220000 | GSE98119 | mouse |  |
| 9 | Chr4: 64180001–64260000<br>Chr4: 64180001–65360000 | GSE98119 | mouse |  |
| 10 | Chr4: 65260001–65340000<br>Chr4: 62800001–65340000 | GSE98119 | mouse |  |
| 11 | Chr9: 76080001–76160000<br>Chr9: 76080001–77260000 | GSE98119 | mouse |  |
| 12 | Chr6: 98450001–98490000<br>Chr6: 98450001–99580000 | GSE82144 | mouse |  |
| 13 | Chr8: 105760001–105840000<br>Chr8: 105760001–106820000 | GSE63525 | human |  |
| 14 | Chr8: 106320001–106360000<br>Chr8: 106320001–106820000 | GSE63525 | human |  |
| 15 | Chr14: 89460001–89520000<br>Chr14: 89460001–90460000 | GSE63525 | human | Gupta et al. [32] |

**Table S1** (continued)

| **N** | **Stripe region** | **GEO/4DN** | **Species** | **Reference** |
| --- | --- | --- | --- | --- |
| 16 | Chr9: 93600001–93660000<br>Chr9: 92860001–93660000 | GSE63525 | human |  |
| 17 | Chr12: 40720001–40800000<br>Chr12: 40200001–40800000 | GSE98119 | mouse |  |
| 18 | Chr12: 41540001–41600000<br>Chr12: 41540001–42280000 | GSE98119 | mouse |  |
| 19 | Chr12: 41820001–41870000<br>Chr12: 41820001–42280000 | GSE98119 | mouse |  |
| 20 | Chr16: 23945001–23990000<br>Chr16: 23945001–25360000 | GSE82144 | mouse | Yoon et al. [31] |
| 21 | Chr16: 25040001–25080000<br>Chr16: 23950001–25080000 | GSE82144 | mouse |  |
| 22 | Chr16: 25240001–25280000<br>Chr16: 23950001–25280000 | GSE82144 | mouse |  |
| 23 | Chr10: 121470001–121505000<br>Chr10: 121050001–121505000 | GSE82144 | mouse |  |
| 24 | Chr1: 130700001–130740000<br>Chr1: 130200001–130740000 | GSE82144 | mouse | Rowley et al. [56] |
| 25 | Chr2: 223160001–223200000<br>Chr2: 223160001–223500000 | GSE63525 | human |  |
| 26 | Chr1: 75500001–75560000<br>Chr1: 74900001–75560000 | GSE116794(Inv1) | mouse |  |
| 27 | Chr1: 75340001–75420000<br>Chr1: 74900001–75420000 | GSE116794(Inv2) | mouse |  |
| 28 | Chr1: 75130001–75210000<br>Chr1: 74800001–75210000 | GSE116794(Inv3) | mouse | Kraft et al. [3] |
| 29 | Chr1: 75020001–75080000<br>Chr1: 74440001–75080000 | GSE116794(Inv4) | mouse |  |
| 30 | Chr1: 132220001–132280000<br>Chr1: 131300001–132280000 | GSE129997(earlyG1) | mouse |  |
| 31 | Chr2: 13220001–13280000<br>Chr2: 13220001–14160000 | GSE129997(earlyG1) | mouse |  |
| 32 | Chr10: 118280001–118320000<br>Chr10: 118280001–118700000 | GSE129997(earlyG1) | mouse |  |
| 33 | Chr1: 132220001–132280000<br>Chr1: 130720001–132280000 | GSE129997(midG1) | mouse |  |
| 34 | Chr2: 13220001–13280000<br>Chr2: 13220001–14300000 | GSE129997(midG1) | mouse | Zhang et al. [15] |
| 35 | Chr10: 118280001–118320000<br>Chr10: 118280001–118700000 | GSE129997(midG1) | mouse |  |

**Table S1** (continued)

| **N** | **Stripe region** | **GEO/4DN** | **Species** | **Reference** |
| --- | --- | --- | --- | --- |
| 36 | Chr1: 132220001–132280000<br>Chr1: 130720001–132280000 | GSE129997(lateG1) | mouse |  |
| 37 | Chr2: 13220001–13280000<br>Chr2: 13220001–14300000 | GSE129997(lateG1) | mouse |  |
| 38 | Chr10: 118280001–118320000<br>Chr10: 118280001–118700000 | GSE129997(lateG1) | mouse |  |
| 39 | Chr3: 3160001–3220000<br>Chr3: 3160001–4300000 | 4DNES21D8SP8 | human |  |
| 40 | Chr3: 3950001–4020000<br>Chr3: 3160001–4020000 | 4DNES21D8SP8 | human |  |
| 41 | Chr3: 4240001–4310000<br>Chr3: 3160001–4310000 | 4DNES21D8SP8 | human |  |
| 42 | Chr3: 4590001–4650000<br>Chr3: 4300001–4650000 | 4DNES21D8SP8 | human |  |
| 43 | Chr3: 4300001–4380000<br>Chr3: 4300001–4960000 | 4DNES21D8SP8 | human |  |
| 44 | Chr3: 3160001–3240000<br>Chr3: 3160001–4300000 | 4DNSRV3SKQ8M | human |  |
| 45 | Chr3: 4240001–4300000<br>Chr3: 3160001–4300000 | 4DNSRV3SKQ8M | human |  |
| 46 | Chr7: 41660001–41740000<br>Chr7: 40140001–41740000 | 4DNESWST3UBH | human |  |
| 47 | Chr7: 41920001–42000000<br>Chr7: 41920001–42940000 | 4DNESWST3UBH | human | Krietenstein et al. [57] |
| 48 | Chr8: 68540001–68600000<br>Chr8: 68540001–69660000 | 4DNESWST3UBH | human |  |
| 49 | Chr15: 66140001–66156000<br>Chr15: 66140001–66596000 | 4DNESWST3UBH | human |  |
| 50 | Chr15: 66492001–66508000<br>Chr15: 66132001–66508000 | 4DNESWST3UBH | human |  |
| 51 | Chr7: 41660001–41740000<br>Chr7: 40140001–41740000 | 4DNSRC6ZVYVP | human |  |
| 52 | Chr7: 41920001–42000000<br>Chr7: 41920001–42940000 | 4DNSRC6ZVYVP | human |  |
| 53 | Chr8: 68540001–68600000<br>Chr8: 68540001–69660000 | 4DNSRC6ZVYVP | human |  |
| 54 | Chr15: 66140001–66156000<br>Chr15: 66140001–66596000 | 4DNSRC6ZVYVP | human |  |
| 55 | Chr15: 66492001–66508000<br>Chr15: 66128001–66508000 | 4DNSRC6ZVYVP | human |  |

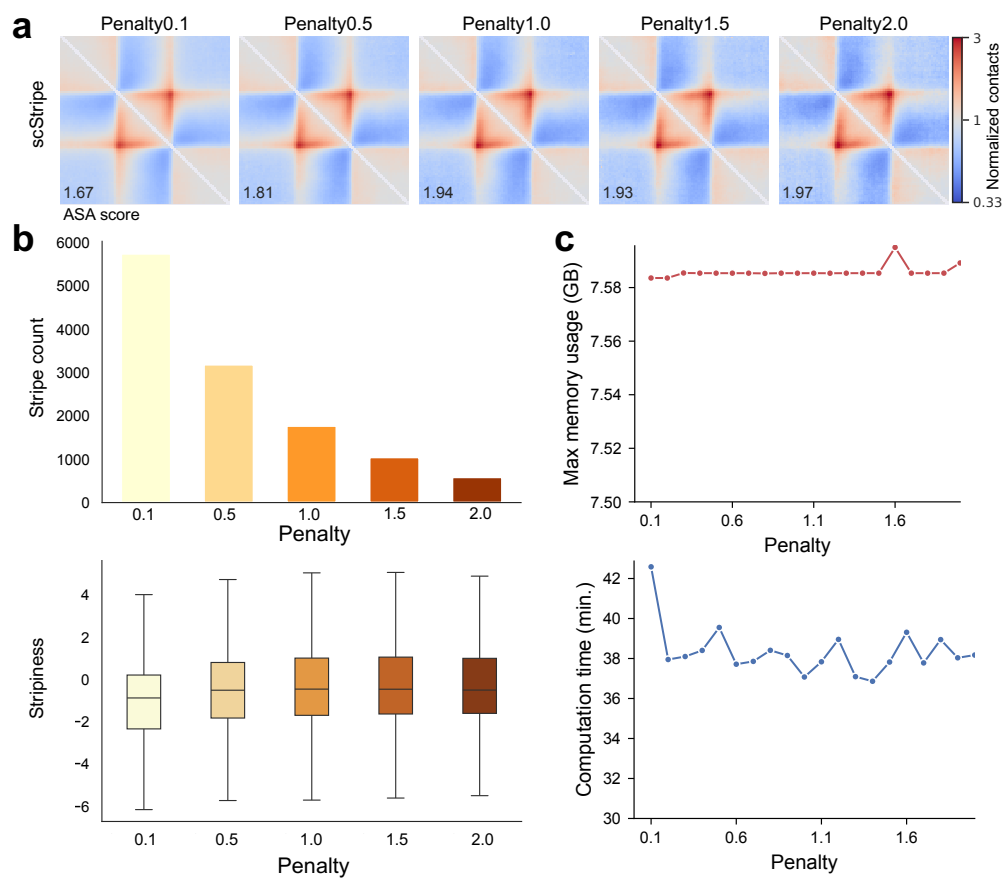

**Figure S1: Sensitivity of scStripe to the change-point penalty.** **a**, Aggregate stripe analysis (ASA) of stripes detected from the excitatory-neuron aggregated scHi-C map across change-point penalty values from 0.1 to 2.0. Change-points were identified along random-matrix-theory–retained eigenvectors by optimizing a global cost function with a penalty term that controls model complexity, such that smaller penalties favor more change-points. ASA increased only modestly as the penalty increased. Numbers in the lower-left corners indicate ASA scores. **b**, Number of detected stripes (top) and stripiness distributions (bottom) across penalty values. Stripe count decreased substantially with increasing penalty, whereas stripiness increased only modestly. **c**, Maximum memory usage (top) and computation time (bottom) across penalty values, showing little variation over the tested range. A penalty of 0.1 was used as the default in subsequent analyses.

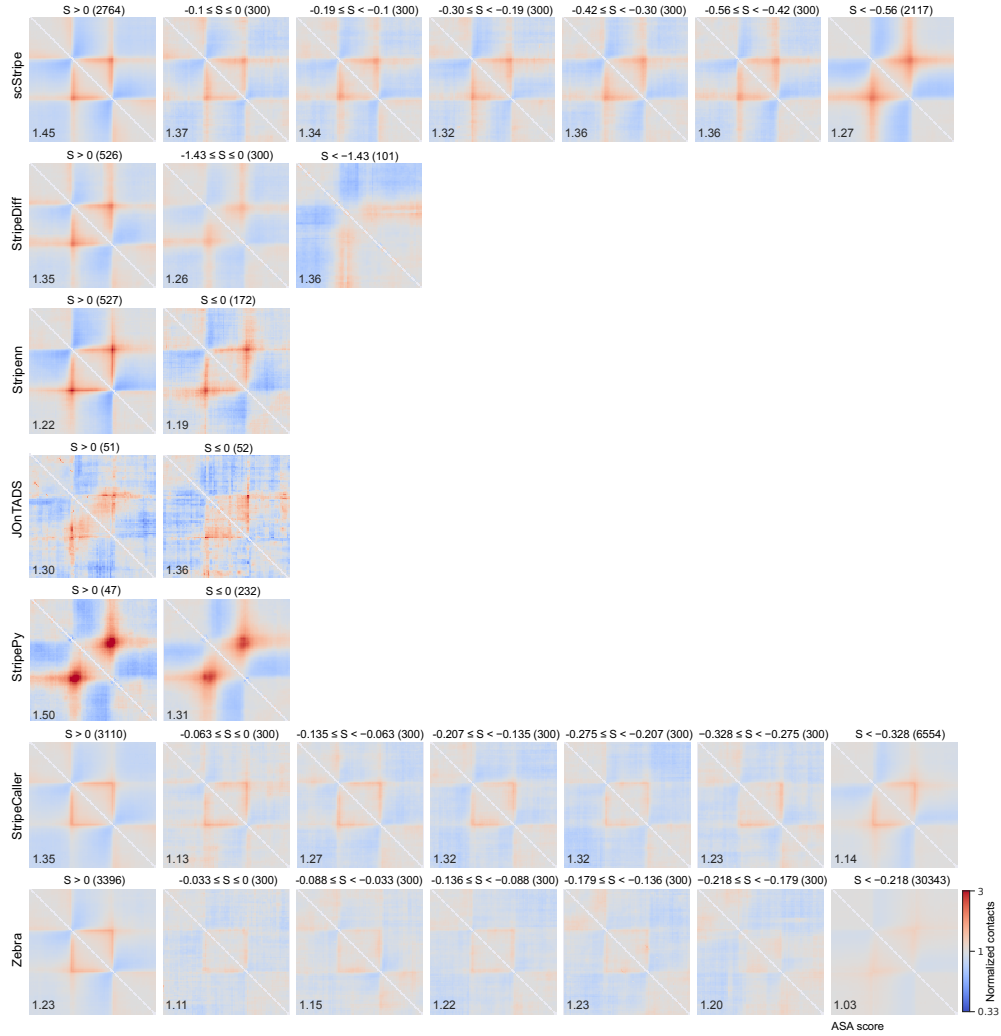

**Figure S2: Aggregate stripe enrichment across stripiness strata and methods.** Stripes were identified from the same bulk NPC Hi-C map by scStripe, StripeDiff, Stripenn, JOnTADS, StripePy, StripeCaller and Zebra and stratified by stripiness ($S$). Following the Stripenn-recommended criterion, all stripes with $S > 0$ were pooled into one group. Stripes with $S \leq 0$ were ranked by $S$ and divided successively into non-overlapping groups of 300 stripes, with the remaining lowest-$S$ stripes pooled into the final group; methods with fewer stripes therefore yielded fewer groups. Each row represents one method, and each panel shows the ASA heatmap for one stripiness stratum. Numbers in the lower-left corners indicate ASA scores. Lower stripiness generally corresponded to weaker aggregate enrichment, but groups with $S \leq 0$ frequently retained clear stripe-centered signal. For scStripe, even the lowest-stripiness group retained substantial aggregate enrichment (ASA 1.27 versus 1.45 for $S > 0$), indicating that a strict $S > 0$ cutoff would exclude stripes supported by aggregate contact enrichment. Accordingly, stripiness was not applied as an additional filter to scStripe calls in the main analyses.

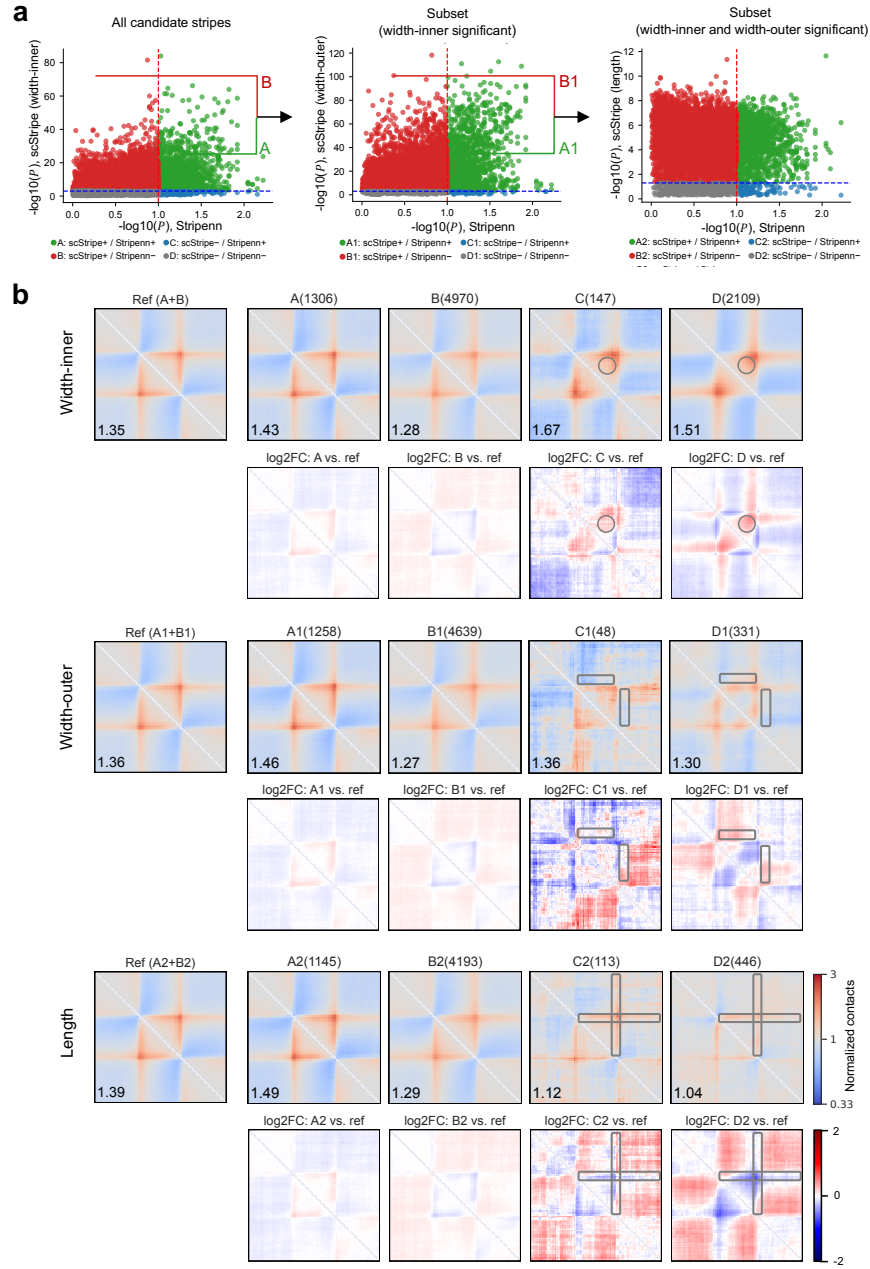

**Figure S3: Stepwise comparison of scStripe test-specific $P$ values with Stripenn.** **a**, Candidate stripes detected by scStripe from bulk NPC Hi-C were evaluated using three one-sided paired $t$-tests, corresponding to the width-inner, width-outer and length comparisons. The two flanking regions were defined relative to stripe orientation as the inner and outer sides. Scatter plots compare $-\log_{10}(P)$ from each scStripe test with the corresponding Stripenn $P$ values; dashed lines indicate the nominal threshold of $P = 0.1$. At the width-inner step, candidate stripes were classified as significant by both methods (A), by scStripe only (B), by Stripenn only (C), or by neither method (D). Stripes significant by scStripe in the width-inner test (A and B) were carried forward to the width-outer test, yielding groups A1–D1. Stripes significant by scStripe in both width tests (A1 and B1) were then evaluated by the length test, yielding groups A2–D2. Thus, only A2 and B2 represent stripes significant by scStripe after all three tests. **b**, Aggregate stripe analysis (ASA) for representative groups in **a**. Stripes in each group were aligned and aggregated to visualize the average stripe-centered contact pattern, with ASA scores shown in the lower-left corners. At each step, the reference aggregate was constructed from stripes significant by scStripe at that step (A+B, A1+B1 or A2+B2), and group-specific $\log_2$ fold-change maps were calculated relative to the corresponding reference. In the width-inner comparison, groups C and D show elevated signal in the highlighted inner-flank region, consistent with failure of the width-inner test. Together, the two width tests select stripes enriched relative to both matched flanking backgrounds, whereas the length test further requires sustained stripe enrichment toward the distal endpoint.

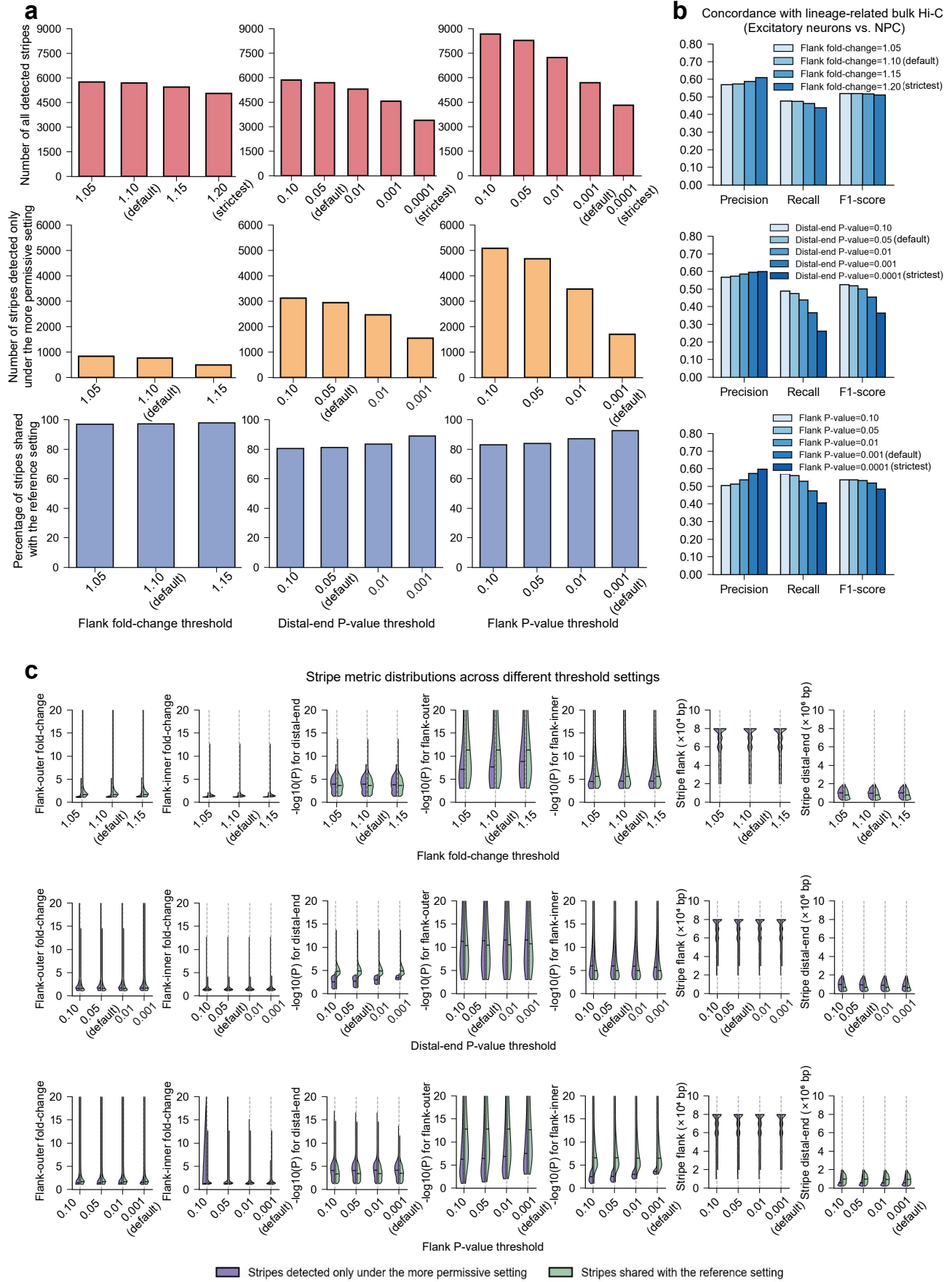

**Figure S4: scStripe-called stripes remain consistent across parameter settings and maintain concordance with lineage-matched bulk Hi-C stripes.** **a**, Numbers of scStripe-called stripes under different parameter settings and their overlap with the reference parameter setting within each parameter series. The numbers of all detected stripes and stripes detected only under the more permissive setting were quantified across flank fold-change thresholds, distal-end $P$-value thresholds and flank $P$-value thresholds. The top row shows the number of all detected stripes, the middle row shows the number of stripes detected only under the more permissive setting, and the bottom row shows the percentage of stripes shared with the reference setting. **b**, Concordance between stripes detected from excitatory-neuron aggregated scHi-C maps and lineage-matched NPC bulk Hi-C stripes across parameter settings. Using the same stripe-matching criterion as in Fig. 2b, Precision, Recall, and F1 score were calculated to quantify concordance between stripes identified by scStripe under each parameter setting and the NPC bulk Hi-C reference set. NPC bulk Hi-C stripes served as the lineage-matched reference and were identified by scStripe from NPC bulk Hi-C using the default parameter setting. This reference set was kept fixed in all comparisons. **c**, Comparison of stripe properties for stripes detected only under the more permissive setting and stripes shared with the reference setting. Distributions of flank-inner fold-change, flank-outer fold-change, $-\log_{10}(P)$ for flank-inner, $-\log_{10}(P)$ for flank-outer, $-\log_{10}(P)$ for distal-end, stripe flank ($\times 10^4$ bp), and stripe distal-end ($\times 10^6$ bp) were compared. The “inner” and “outer” sides were defined in the Hi-C upper-triangle matrix according to stripe orientation. For 3'-stripes, the “inner” side corresponds to the flank region below the stripe and the “outer” side to the flank region above it. For 5'-stripes, the “inner” side corresponds to the left flank and the “outer” side corresponds to the right flank. For each parameter series, the most stringent parameter setting was used as the reference setting and three stripe categories were defined. All detected stripes refer to stripes identified by scStripe under the corresponding parameter setting. Stripes shared with the reference setting denote stripes from a permissive parameter setting whose genomic intervals completely contain at least one stripe identified under the reference parameter setting. Stripes detected only under the more permissive setting denote stripes from a permissive parameter setting whose genomic intervals do not completely contain any stripe from the reference parameter setting.

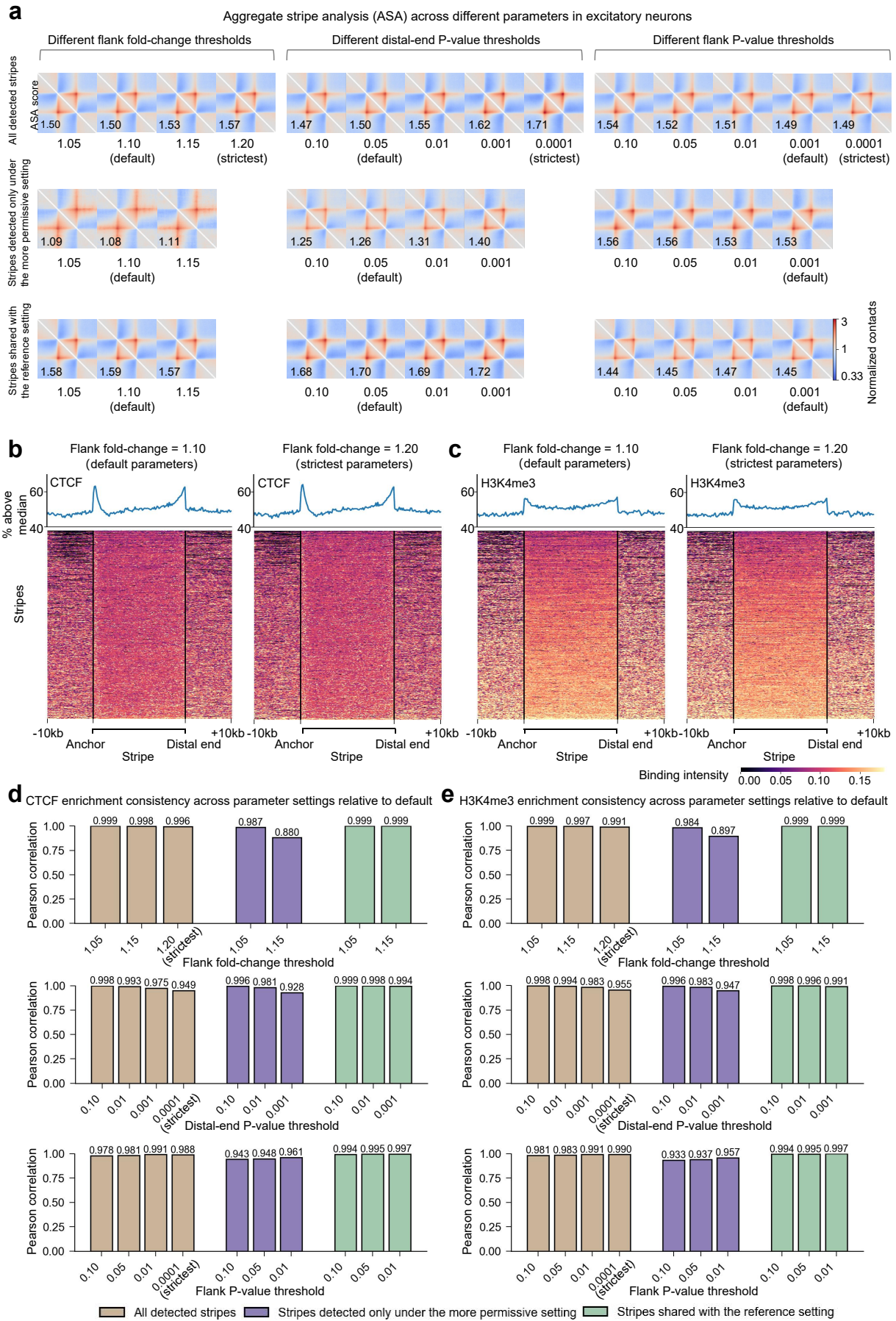

**Figure S5: scStripe-called stripes exhibit stable ASA scores and consistent CTCF and H3K4me3 enrichment across parameter settings.** **a**, Aggregate stripe analysis (ASA) of stripes identified by scStripe under different parameter settings. ASA was performed separately for all detected stripes, stripes detected only under the more permissive setting, and stripes shared with the reference setting. The corresponding ASA score is shown in the lower-left corner of each heatmap. **b**, CTCF enrichment of scStripe-called stripes under the default and reference flank fold-change thresholds. Heatmaps show normalized CTCF ChIP-seq signals aligned to the identified stripes. For each stripe, the stripe body was normalized into 100 bins, with 50 upstream and 50 downstream bins added, yielding a region of 200 bins in total. Vertical black lines mark the stripe anchor and distal end, respectively. Line plots above show, at each relative position, the proportion of stripes with signal greater than the heatmap-wide median, yielding a proportion profile of length 200. **c**, H3K4me3 enrichment of stripes identified by scStripe under the default and reference flank fold-change thresholds, shown as normalized heatmaps and proportion profiles as in **b**. **d**, Consistency of CTCF enrichment across parameter settings relative to the default setting. For each parameter setting and stripe category shown in **a**, a CTCF proportion profile of length 200 was computed as described in **b**. Pearson correlation coefficients were then calculated between each proportion profile and the corresponding profile at the default threshold within the same parameter series. **e**, Consistency of H3K4me3 enrichment across parameter settings relative to the default setting, analyzed as in **d**. CTCF and H3K4me3 signals were processed using the same data sources, normalization procedures, and stripe-rescaling strategy as described in S10d,e. ASA was performed using the same workflow and parameter settings as in Fig. 2c. All stripe sets were identified by scStripe from the excitatory-neuron aggregated scHi-C map.

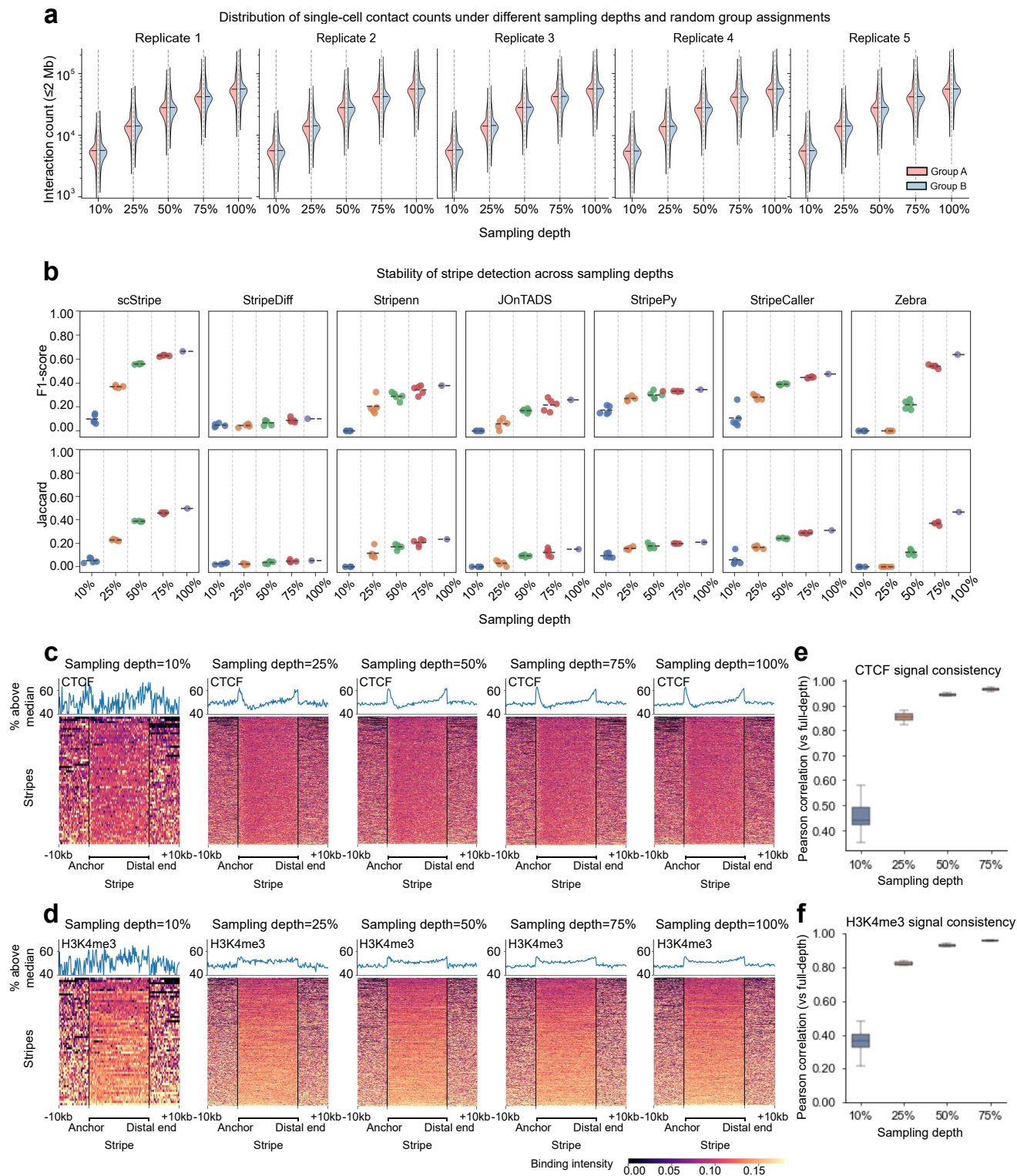

**Figure S6: scStripe maintains stable stripe-calling concordance across sampling depths, with consistent CTCF and H3K4me3 enrichment patterns.** **a**, Distributions of single-cell contact counts across sampling depths. All excitatory neurons were downsampled without replacement, and five independent replicates were generated at each sampling depth of 10%, 25%, 50% and 75%. The full-depth condition (100%) corresponded to the raw single-cell contact data. Cells were subsequently randomly partitioned into two equal-sized, non-overlapping groups (A and B). Violin plots show the distributions of cis-contact counts within a genomic distance $\leq 2\text{ Mb}$ for cells in groups A and B across the five replicates. **b**, Comparison of stripe-calling stability across sampling depths for different methods in downsampled excitatory-neuron aggregated scHi-C maps. For

**Figure S6:** (Continued.) each sampling depth and replicate, aggregated scHi-C maps were generated separately from cells in groups A and B, followed by independent stripe calling. To evaluate stripe-calling stability across sampling depths, concordance between stripe sets identified from groups A and B was assessed using the F1 score and Jaccard index. Results are shown for scStripe, StripeDiff, Stripenn, JOnTADS, StripePy, StripeCaller, and Zebra across sampling depths. scStripe achieved the highest or near-highest F1 scores and Jaccard indices under most sampling depths and maintained stable performance at sampling depths of 25% and above. **c**, CTCF enrichment of scStripe-called stripes across sampling depths. Heatmaps show normalized CTCF ChIP-seq signals aligned to rescaled stripe regions across sampling depths for a representative down-sampling replicate (replicate 1). For each stripe, the stripe body was normalized to 100 bins, with 50 upstream and 50 downstream bins added, yielding a region of 200 bins in total. Vertical black lines mark the stripe anchor and distal end, respectively. Line plots above show, at each relative position, the proportion of stripes with signal greater than the heatmap-wide median, yielding a proportion profile of length 200. **d**, H3K4me3 enrichment of scStripe-called stripes across sampling depths, shown as normalized heatmaps and proportion profiles as in **c**. **e**, Consistency of CTCF enrichment across sampling depths relative to the full-depth condition. For each stripe set identified under a downsampled condition, a CTCF proportion profile of length 200 was computed as described in **c**. Pearson correlation coefficients were calculated between the proportion profile from each downsampled condition and the corresponding full-depth profile within the same replicate, and were then summarized across five independent downsampling replicates for each sampling depth. **f**, Consistency of H3K4me3 enrichment across sampling depths relative to the full-depth condition, analyzed as in **e**. Downsampling was performed independently for each cell by random sampling without replacement, and the A/B cell partition was kept fixed throughout all analyses. CTCF and H3K4me3 signals were processed using the same data sources, normalization procedures, stripe-rescaling strategy and analysis workflow as in Fig. S10d,e. Stripe sets in panels **c–f** were identified by scStripe from the corresponding downsampled aggregated scHi-C maps.

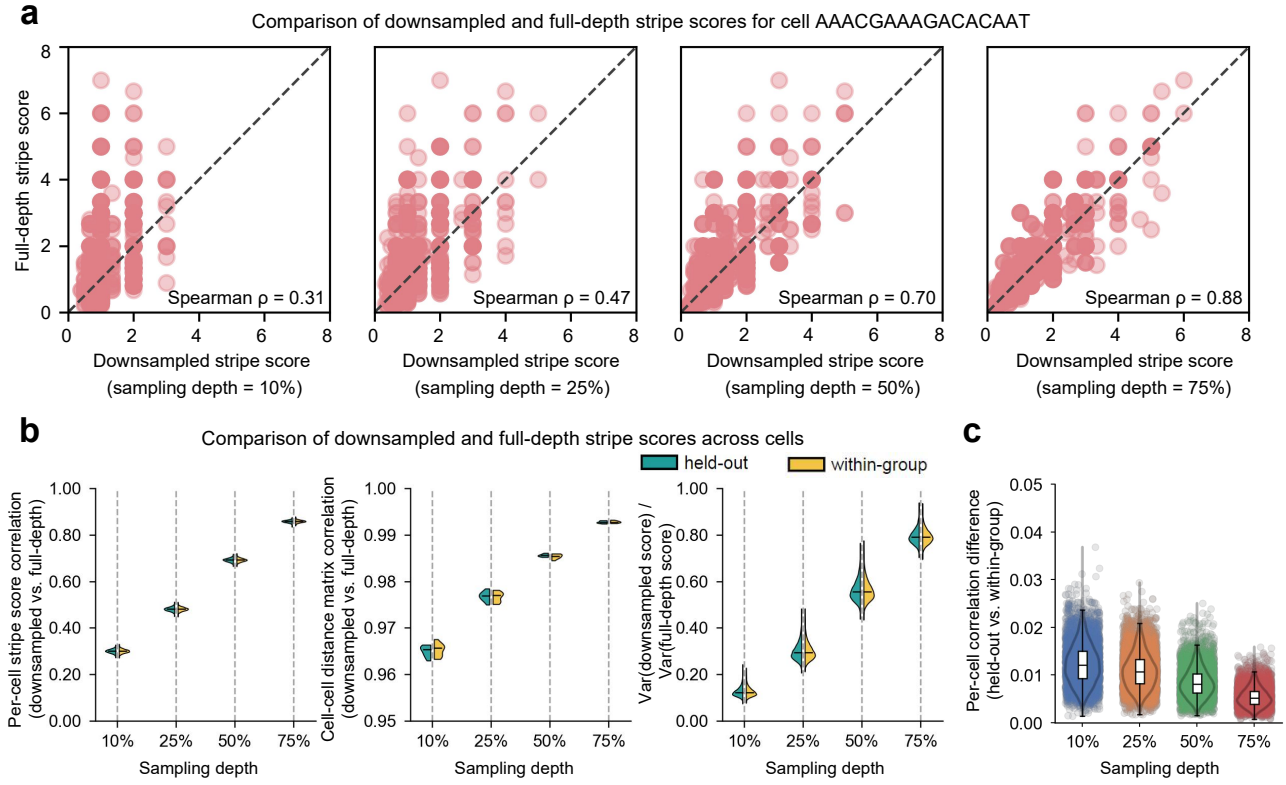

**Figure S7: Per-cell stripe scores computed by scStripe remain stable across sampling depths and show limited sensitivity to self-inclusion.** **a**, Consistency of per-cell stripe scores across sampling depths for a representative single cell. Using cell AAACGAAAGACACAAT as an example, stripe scores obtained at sampling depths of 10%, 25%, 50% and 75% in replicate 1 were compared with those from the full-depth condition. Each point represents one stripe identified by scStripe from the full-depth aggregated scHi-C map generated from group A using raw single-cell contact counts. The x-axis shows stripe scores under the downsampled condition, and the y-axis shows stripe scores under the corresponding full-depth condition. The corresponding Spearman correlation coefficient is shown in the lower-right corner. **b**, Consistency of per-cell stripe scores across sampling depths. The left panel shows Spearman correlations between per-cell stripe-score vectors from downsampled and full-depth conditions, averaged across five downsampling replicates. The middle panel shows the consistency of cell-cell distance matrices between downsampled and full-depth conditions. For each condition, a cell × stripe score matrix was obtained from stripe scores of all stripes across all cells. Stripe score vectors were then used to compute pairwise Euclidean distance matrices between cells. The upper triangular elements, excluding the diagonal, of the cell-cell distance matrices from the downsampled and full-depth conditions were extracted, yielding two vectors of pairwise cell-cell distances, and the Spearman correlation coefficient between these vectors was calculated. The right panel shows changes in stripe-score variability across cells relative to the full-depth condition. For each stripe, the ratio of stripe score variance across cells between the downsampled and full-depth conditions was calculated and averaged across five downsampling replicates. All three metrics were evaluated separately for within-group scoring and held-out scoring. **c**, Difference between held-out scoring and within-group scoring. For each cell, the Spearman correlation coefficient between stripe score vectors from the downsampled and full-depth conditions was first calculated. The absolute difference between the corresponding correlation coefficients obtained from held-out scoring and within-group scoring was then computed. Results were averaged across five downsampling replicates and summarized by sampling depth. Held-out scoring and within-group scoring were both performed using fixed stripe sets identified by scStripe from aggregated scHi-C maps generated without downsampling. Specifically, aggregated maps were constructed separately for the two non-overlapping groups A and B from raw single-cell contact counts, yielding stripe sets $S_A$ and $S_B$, respectively. Stripe scores were subsequently calculated from single-cell contact data across all sampling depths and downsampling replicates. Held-out scoring refers to calculating stripe scores for stripes in $S_A$ using cells from group B, or for stripes in $S_B$ using cells from group A. Within-group scoring refers to calculating stripe scores for stripes in $S_A$ using cells from group A, or for stripes in $S_B$ using cells from group B.

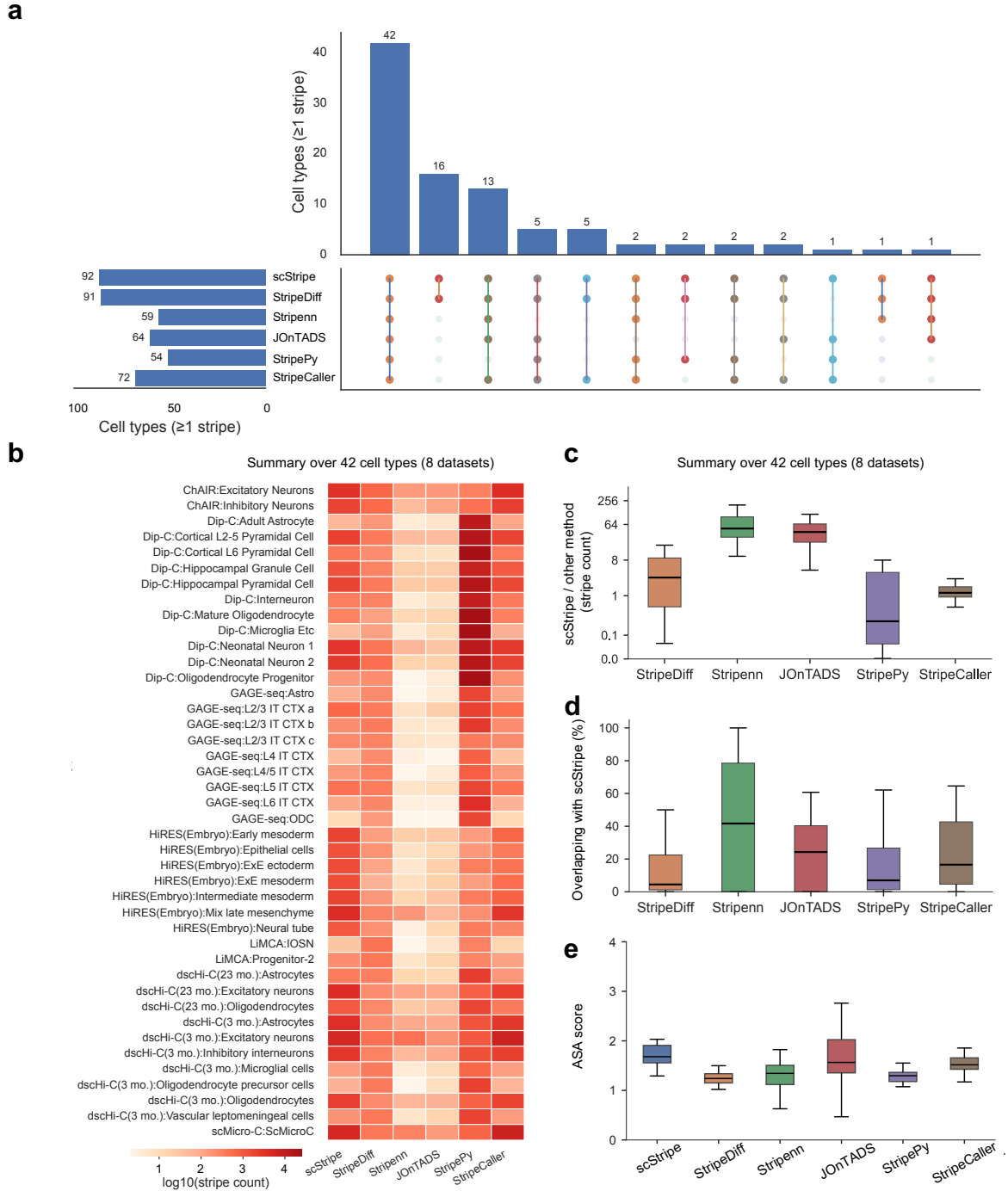

**Figure S8: Cross-cell-type benchmarking of stripe detection from aggregated scHi-C maps.** **a**, UpSet plot summarizing stripe-detection breadth across 94 cell types. Horizontal bars show the number of cell types in which each method detected at least one stripe. Vertical bars show the number of cell types corresponding to each combination of methods, with the dot matrix indicating method membership. The 42 cell types across eight datasets in which all six callers detected at least one stripe were retained for the cross-cell-type comparisons in **b–e**. **b**, Heatmap showing the number of stripes detected by each method across the 42 cell types. Stripe counts are shown on a $\log_{10}$ scale. **c**, Distribution of stripe-count ratios between scStripe and each alternative caller across the 42 cell types. The y-axis shows the ratio of stripe counts (scStripe/alternative caller) on a $\log_2$ scale. Ratios greater than 1 indicate more stripes detected by scStripe, whereas ratios below 1 indicate more stripes detected by the alternative caller. **d**, Fraction of stripes detected by each alternative caller that overlap scStripe calls across the 42 cell types. **e**, ASA scores for stripes detected by each method across the 42 cell types, quantifying aggregate stripe-centered enrichment relative to matched flanking regions.

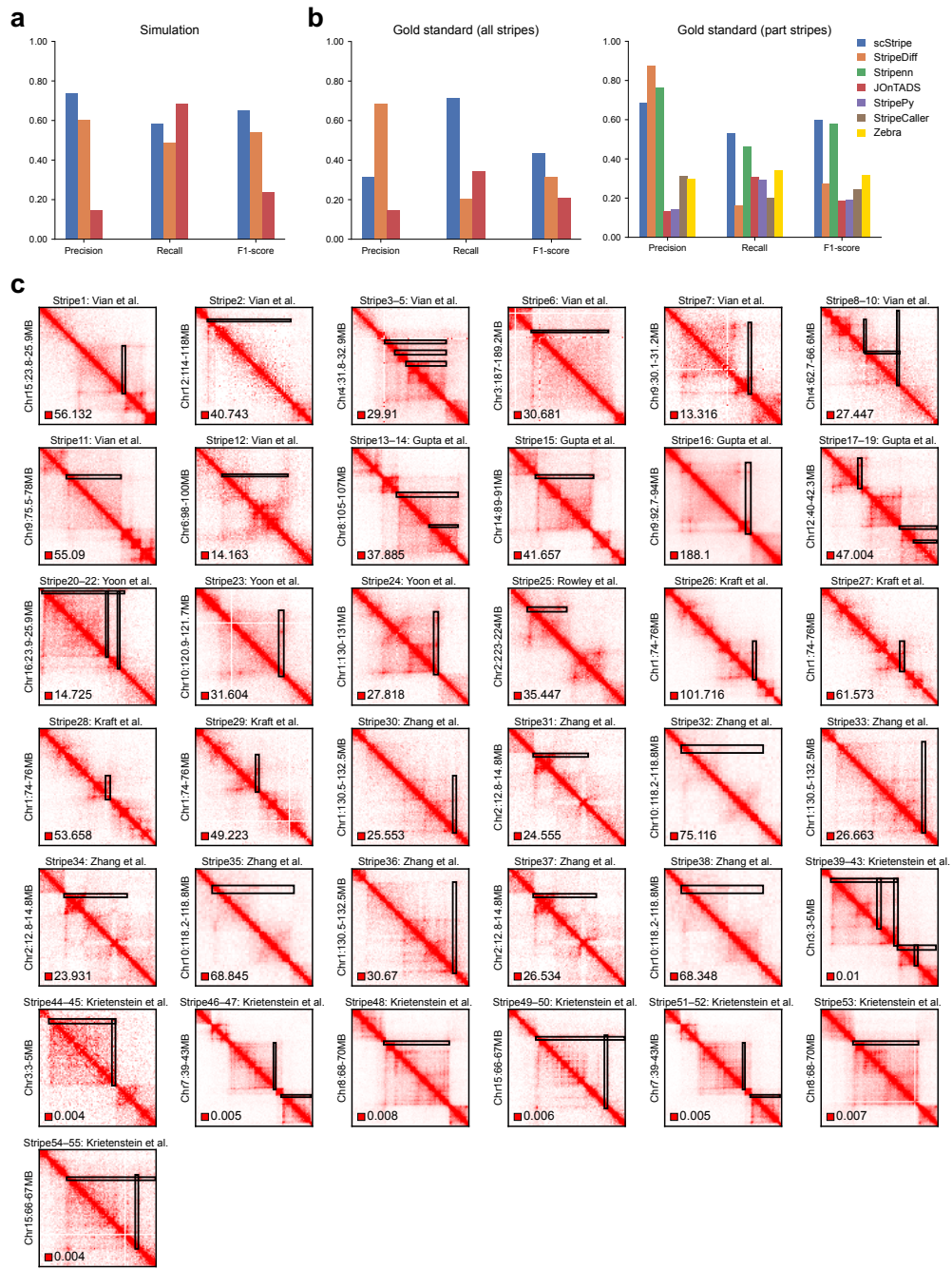

**Figure S9: Benchmarking on simulated contact maps and literature-reported bulk Hi-C stripes.** **a**, Performance on 100 simulated Hi-C contact maps generated with SimuStripe [32]. scStripe, StripeDiff and JOnTADS, which can operate directly on the simulated contact matrices, were evaluated. Stripenn, StripePy, StripeCaller and Zebra require Cooler-format or whole-chromosome inputs and were therefore not included. Precision, recall and F1 score were calculated for each simulation and summarized across the 100 simulations. **b**, Benchmarking against a manually curated set of 55 stripes reported in seven bulk Hi-C studies (Methods). *Left*, results for all 55 literature-reported stripes using scStripe, StripeDiff and JOnTADS, which do not require Cooler-format input. *Right*, results for the subset of 51 stripes for which the corresponding bulk Hi-C data were available in Cooler format, allowing Stripenn, StripePy, StripeCaller and Zebra to be run and all seven methods to be compared. Sources and genomic coordinates of the curated stripes are provided in Supplementary Table S1. **c**, Visualization of the 55 literature-reported stripes in their corresponding bulk Hi-C maps. Black boxes indicate the stripes reported in the original publications.

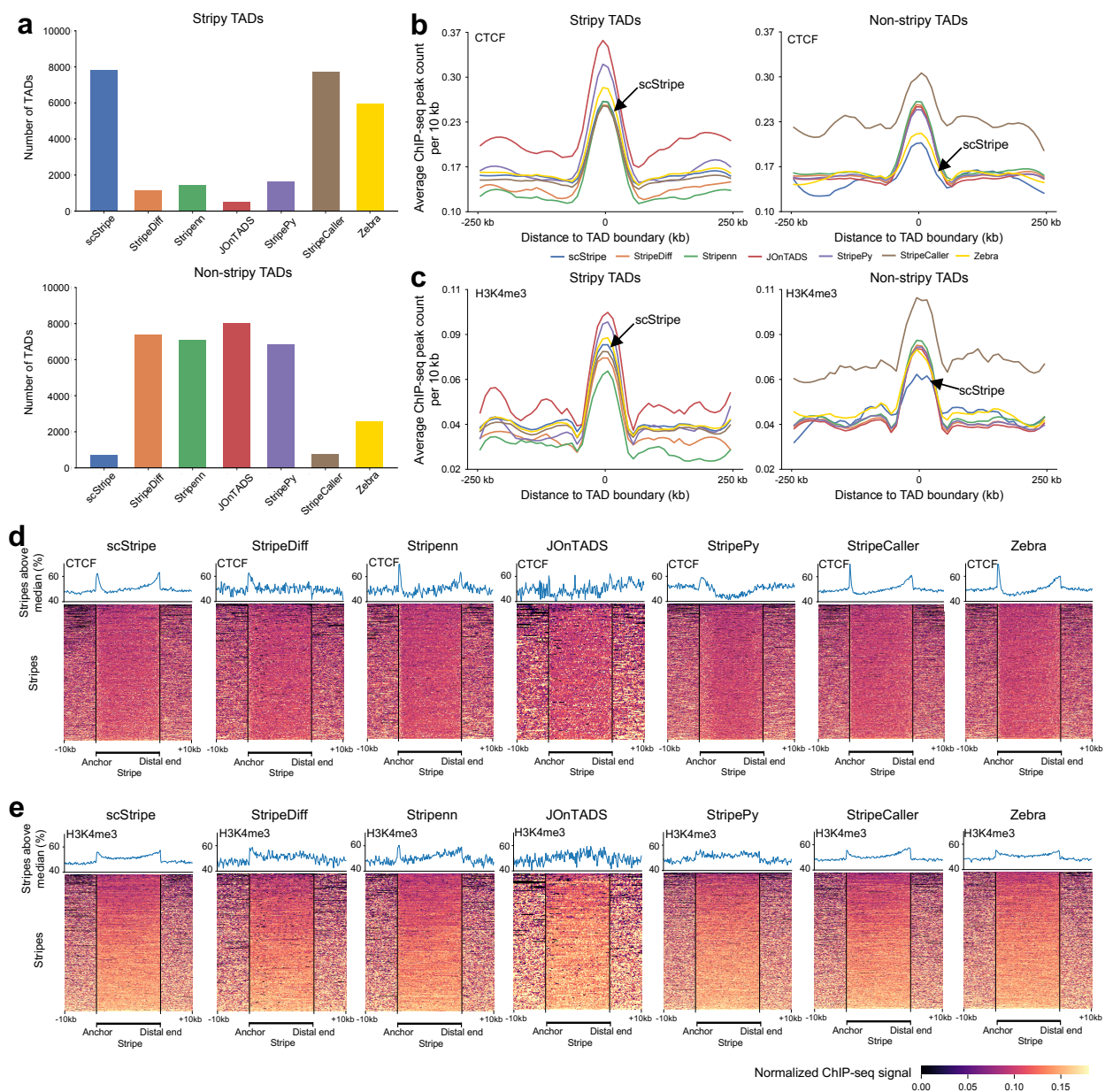

**Figure S10: Stripes detected from aggregated scHi-C maps show CTCF and H3K4me3 enrichment at stripy TAD boundaries and along stripe regions.** **a**, Numbers of stripy and non-stripy TADs associated with the stripe calls from each method. For each caller, independently called TADs were classified as stripy if they overlapped at least one detected stripe and as non-stripy otherwise [31]. **b**, Average CTCF ChIP-seq peak counts per 10 kb centered on TAD boundaries, shown separately for stripy TADs (*left*) and non-stripy TADs (*right*) according to the stripe calls from each method. **c**, Average H3K4me3 ChIP-seq peak counts per 10 kb centered on the same TAD boundaries. **d**, Normalized CTCF ChIP-seq signal aligned to stripe coordinates for each caller. Stripe regions between the anchor and distal end were rescaled to a common length, with 10 kb flanking regions retained on both sides. Vertical black lines mark the stripe anchor and distal end. Line plots above the heatmaps show, at each relative position, the proportion of stripes with signal above the heatmap-wide median. **e**, Normalized H3K4me3 ChIP-seq signal aligned and summarized as in **d**. All panels use stripes and TADs derived at 10 kb resolution from the excitatory-neuron aggregated scHi-C map. CTCF and H3K4me3 signals were obtained from lineage-related NPC ChIP-seq data [41]. The stripe anchor denotes the genomic locus from which the directional interaction trajectory originates, and the distal end denotes the terminal extent of the stripe [2].

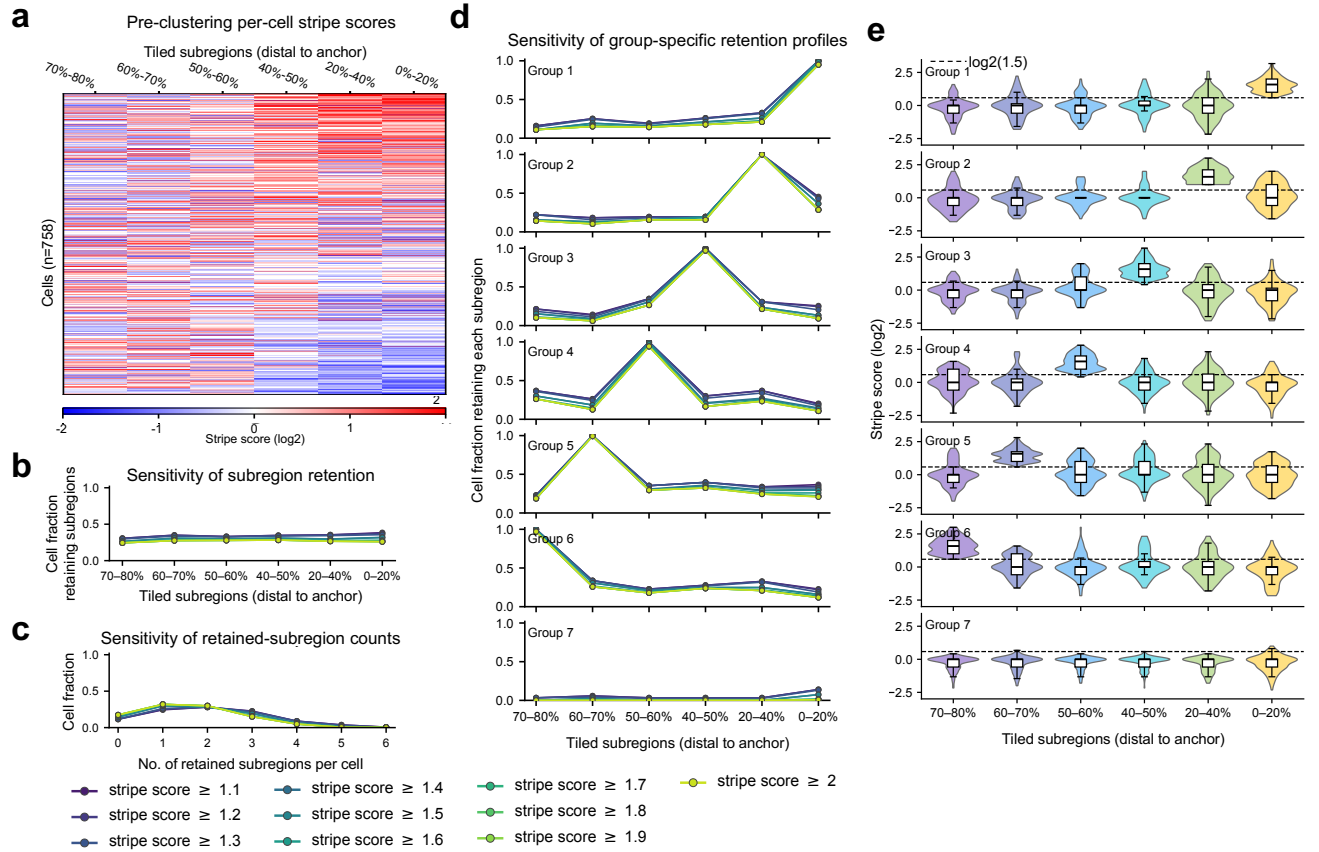

**Figure S11: Per-cell *EBF1* stripe heterogeneity and robustness across stripe-score thresholds.** **a**, Heatmap of the six tiled per-cell stripe scores shown in Fig. 4e, with cells reordered using the per-cell stripe score calculated from the distal 50% of the stripe body. The tiled subregions are ordered from distal to anchor. This ordering visualizes cell-to-cell variation without using the hierarchical-clustering group labels. **b**, Fraction of cells retaining each tiled subregion across stripe-score thresholds from 1.1 to 2.0. **c**, Distribution of the number of retained subregions per cell across the same thresholds. **d**, Group-specific fractions of cells retaining each tiled subregion across the same thresholds. Groups correspond to those defined in Fig. 4e. **e**, Violin plots of $\log_2$ per-cell stripe scores across the six tiled subregions for Groups 1–7. The dashed line marks $\log_2(1.5)$.

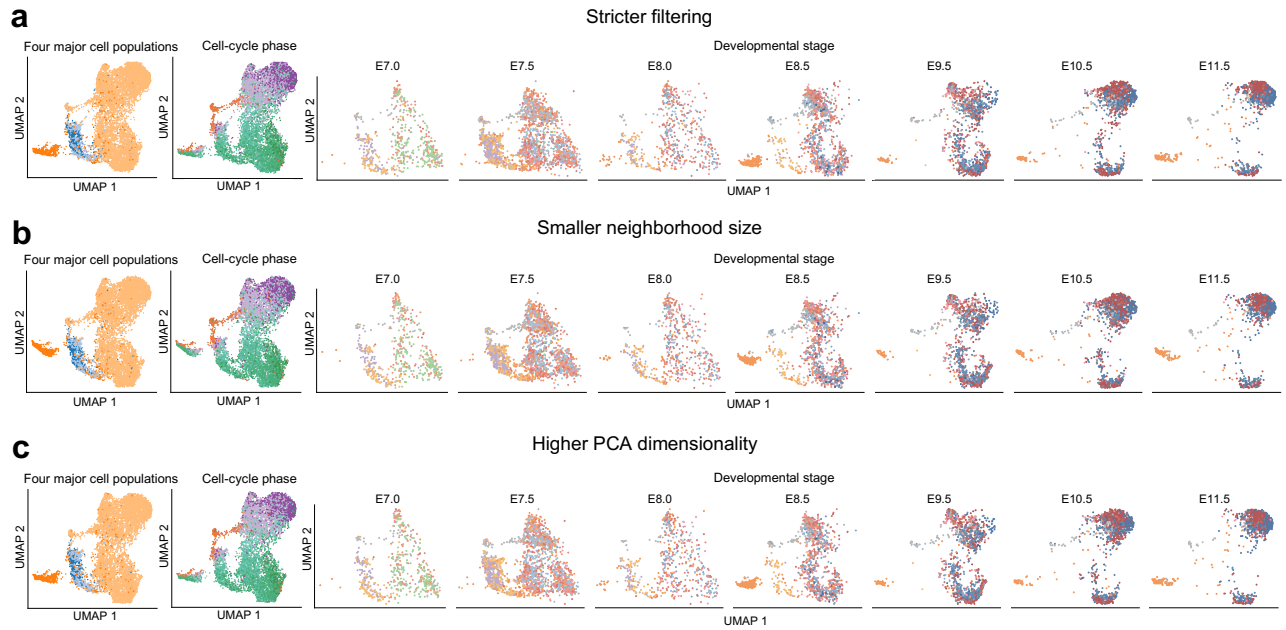

**Figure S12: Robustness of genome-wide per-cell stripe-score embeddings to analysis parameters.** UMAP embeddings based solely on genome-wide per-cell stripe scores after modifying one analysis setting relative to the default setting used in Fig. 5b. In each panel, the embedding is shown by four major cell populations and cell-cycle phase, followed by stage-specific views spanning E7.0–E11.5, with annotation colors following Fig. 5b. **a**, Stricter feature filtering with `min_features = 5000` and `min_cells = 500`. **b**, Smaller neighborhood size with `n_neighbors = 25`. **c**, Higher PCA dimensionality with `n_pca = 30`. The default setting in Fig. 5b was `min_features = 3000`, `min_cells = 300`, `n_neighbors = 35`, `n_pca = 20` and `resolution = 1.0`. Per-cell stripe scores were computed using the same genome-wide profiling mode as in Fig. 5. All analyses use the HiRES dataset [24].

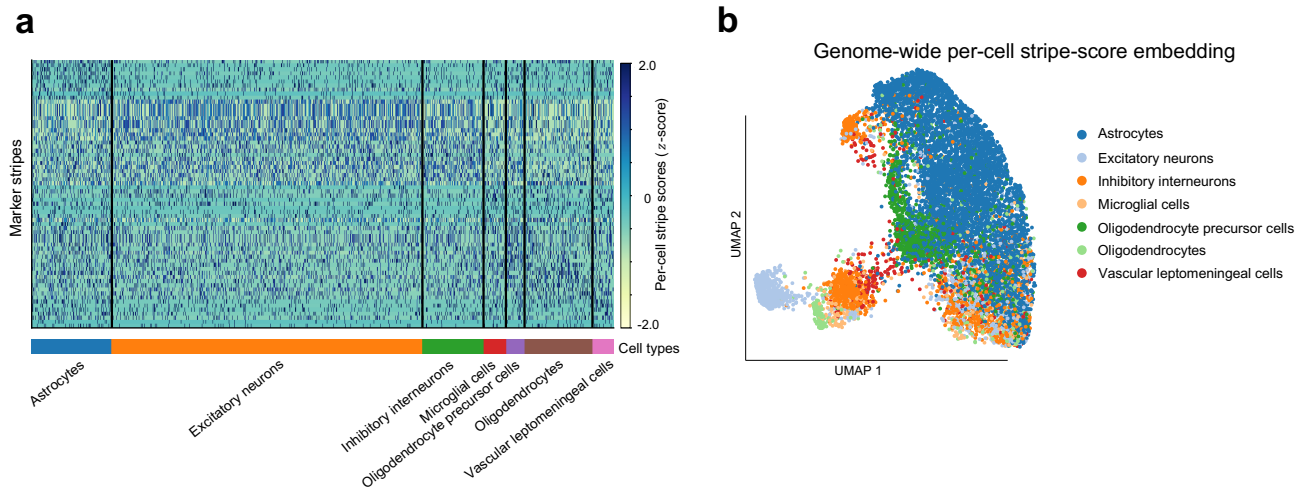

**Figure S13: Genome-wide per-cell stripe scores capture cell-type-associated variation in an independent dscHi-C dataset.** **a**, Heatmap of $z$-scored per-cell stripe scores for cell-type marker stripes, with cells grouped by seven annotated brain cell types. **b**, UMAP embedding based solely on genome-wide per-cell stripe scores, colored by the same seven cell types. For genome-wide profiling, each stripe was represented by one score per cell using the distal 50% of the stripe body relative to matched flanking regions. All analyses use the dscHi-C dataset [22], with cell-type annotations following the original study.

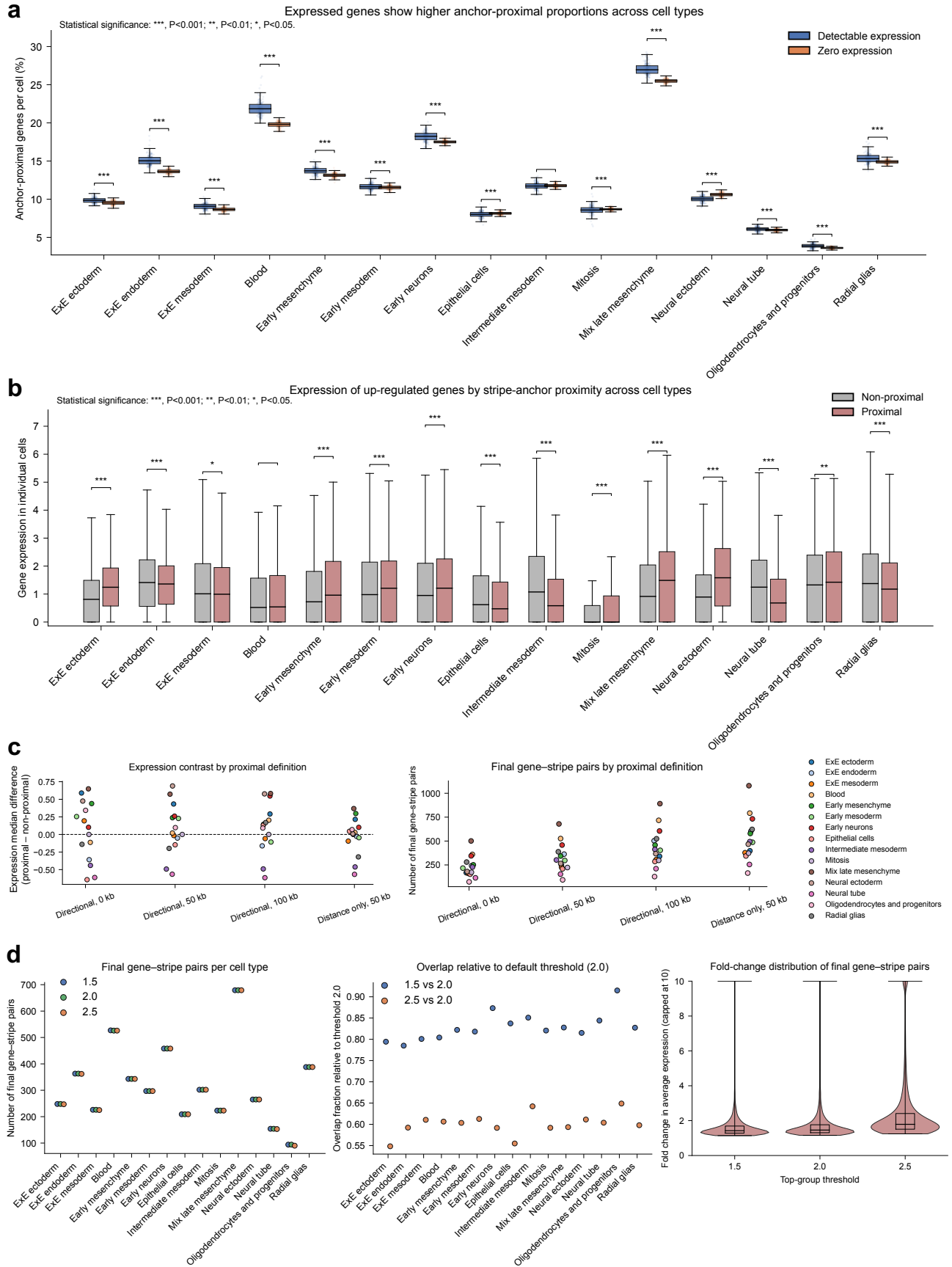

**Figure S14: Cross-cell-type consistency and robustness of stripe-expression relationships in the HiRES dataset.** a, Across cell types, distributions of the per-cell proportion of anchor-proximal genes

**Figure S14:** (Continued.) among genes with detectable or zero expression. Statistical significance is indicated above each comparison. **b**, Across cell types, expression distributions for the union of the top 100 up-regulated genes from each cell type, stratified by whether genes were proximal or non-proximal to stripe anchors. Statistical significance is indicated above each comparison. **c**, Robustness to the definition of stripe-anchor proximity. Left, median expression differences between proximal and non-proximal genes under four definitions: Directional, 0 kb; Directional, 50 kb; Directional, 100 kb; and Distance only, 50 kb. Under the directional definitions, gene TSSs were required to lie within the directional span of the stripe and within the indicated distance from the stripe anchor; the distance-only definition did not impose the directional-span requirement. Right, numbers of final gene–stripe pairs identified under the same four definitions. **d**, Robustness to the stripe-score threshold used to define the top group in gene–stripe pair analysis. Left, numbers of final gene–stripe pairs per cell type at stripe-score thresholds of 1.5, 2.0 and 2.5. Middle, overlap fractions of final gene–stripe pairs identified at thresholds of 1.5 and 2.5 relative to those identified at the default threshold of 2.0. Right, distributions of fold changes in average gene expression between the top and bottom groups for final gene–stripe pairs under each threshold. Final gene–stripe pairs were obtained by ranking candidate proximal gene–stripe pairs by fold change in average gene expression between the top and bottom groups and retaining the top 25% within each cell type. All analyses used the HiRES dataset [24] analyzed in Fig. 6, with stripes identified from aggregated scHi-C maps for each cell type and per-cell stripe scores computed at 10 kb resolution.

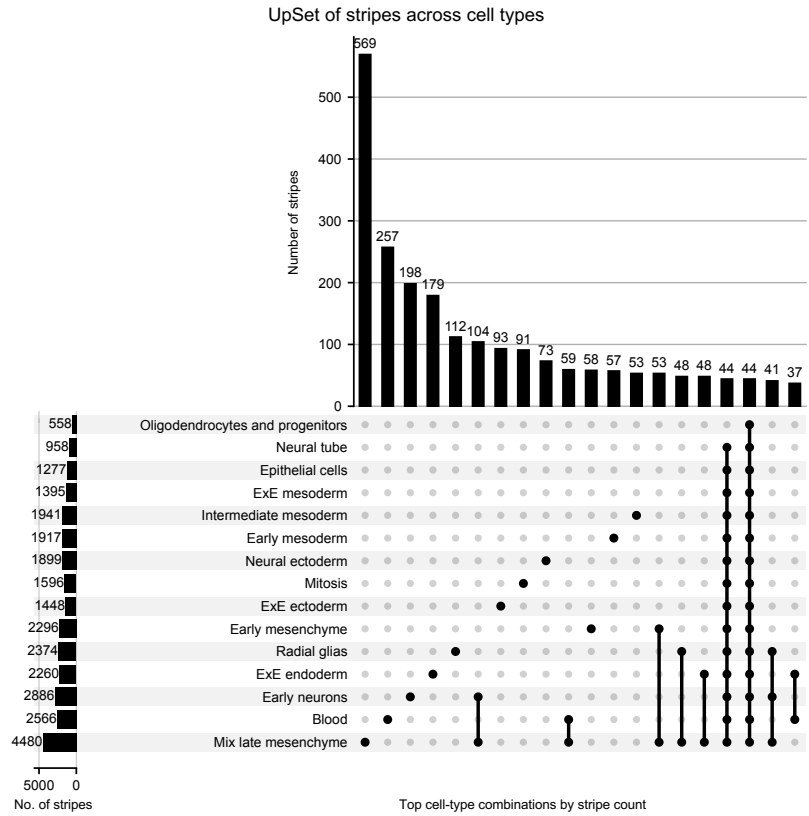

**Figure S15: Cell-type sharing of stripes across the HiRES cell types.** UpSet plot showing unique and shared stripes across cell types. Vertical bars show stripe counts for the leading cell-type combinations, with the dot matrix indicating cell-type membership, and horizontal bars show the total number of stripes detected in each cell type. The leading combinations were predominantly restricted to a single cell type. Stripes were identified from aggregated scHi-C maps for each cell type in the HiRES dataset [24] used in Fig. 6.

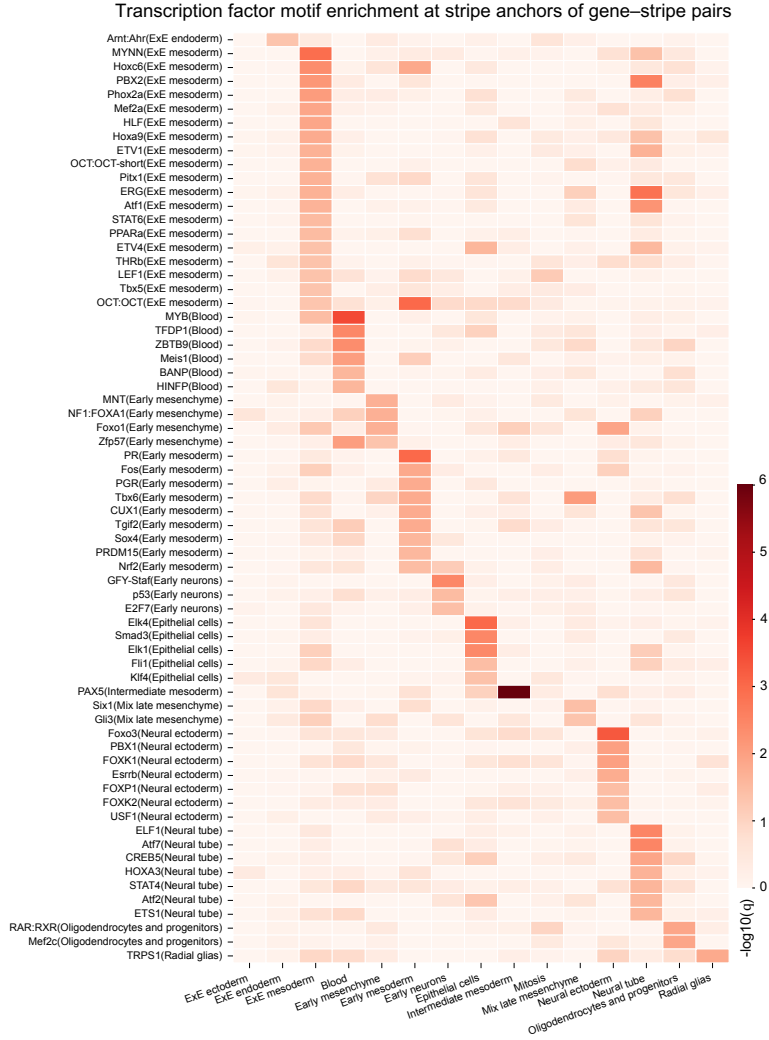

**Figure S16: Transcription factor motif enrichment at stripe anchors of gene–stripe pairs across cell types.** Heatmap of significantly enriched transcription factor motifs at stripe anchors of gene–stripe pairs across cell types. Heatmap values show $-\log_{10}(q)$; only significantly enriched motifs are shown. All analyses used the HiRES dataset [24] analyzed in Fig. 6.
